# Structure-based identification of GIRK2-PIP2 modulators among known drugs and metabolites using docking, MM-GBSA, ADMET, and molecular dynamics

**DOI:** 10.1101/2025.05.13.653795

**Authors:** Danko Jeremic, Lydia Jiménez-Díaz, Juan D. Navarro-López

## Abstract

G protein-gated inwardly rectifying potassium (GIRK) channels are key regulators of neuronal excitability, making them promising therapeutic targets for central nervous system disorders. Their activation depends on phosphatidylinositol-4,5-bisphosphate (PIP_2_), which stabilizes the channel’s open state. A deeper understanding of GIRK-PIP_2_ interactions could uncover new physiological roles and pave the way for therapies that modulate channel function.

This study aimed to advance the targeting of GIRK channels at the PIP_2_-binding site. Over one million compounds were screened against GIRK2 (PDB ID: 4KFM) using high-throughput virtual screening. A core constraint with a root-mean-square deviation (RMSD) < 2 Å was applied to assure the accuracy and binding close to PIP_2_-binding site. The top-scoring ligands were redocked with Glide (SP, XP) and binding free energy was estimated using Molecular Mechanics Generalized Born Surface Area method. The most promising compounds were analyzed for pharmacokinetic/physicochemical properties, followed by molecular dynamics (MD) simulations over 200 ns in membrane bilayer.

MD analysis revealed three known compounds (Rosuvastatin, CID: 54365126 and 7304563) as potential competitive GIRK2 modulators, exhibiting stable interactions with residues critical for binding endogenous activators (PIP_2_, cholesterol), and GIRK-acting drugs. Docking analyses also revealed strong binding to GIRK2 for various metabolites, including leukotrienes, resolvins, acyl-CoAs, and polyphosphates, including adenosine-triphosphate (ATP) and thiamine-triphosphate.

Notably, some of the identified compounds can affect similar ion channels, indicating potential cross-reactivity with GIRK2. Furthermore, the binding modes of acyl-CoAs and polyphosphates closely resemble PIP_2_’s hydrophobic and phosphate group engagement. Together, these findings offer promising candidates for experimental validation and therapeutic development.

**Graphical abstract:** 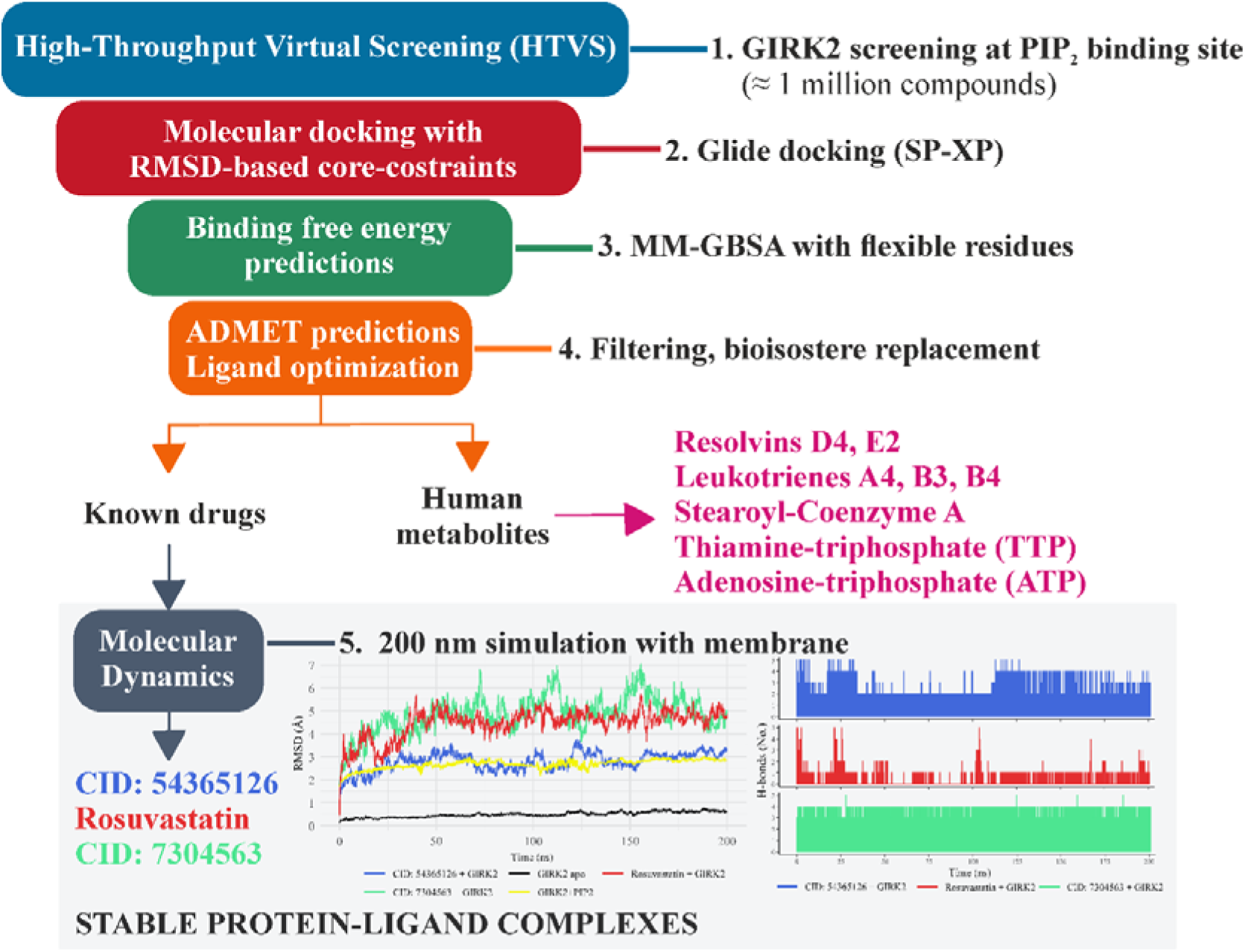

Abbreviations: ADMET – Absorption, Distribution, Metabolism, Elimination, Toxicity; CID – PubChem Compound Identification; MM-GBSA – Molecular Mechanics with Generalized Born and Surface Area solvation; RMSD - Root-Mean-Square Deviation; SP – Standard Precision; PIP_2_ – phosphatidylinositol-4,5-bisphosphate; XP – Extra Precision.

**HIGHLIGHTS:**

- Multi-target screening identifies selective modulators of the GIRK2 channel
- Ligands target the PIP2-binding site, a novel interface for GIRK2 modulation
- MM-GBSA confirms binding affinity and ligand stability post-docking
- Dynamics and bioinformatics predict selectivity and off-target interactions
- Statins and leukotriene-modifying drugs are strong GIRK2 modulator candidates

## 1. INTRODUCTION

G protein-coupled inwardly rectifying potassium (GIRK) channels are a family of lipid-gated inwardly rectifying potassium channels that play a critical role in regulating cellular excitability and signaling in various tissues, particularly in the central nervous system and the heart. These channels are activated by phosphatidylinositol-4,5-bisphosphate (PIP_2_) and modulated by a cascade of molecular signals triggered when neurotransmitters or neuromodulators bind to nearby G protein-coupled receptors (GPCRs). A variety of GPCRs can regulate GIRK channels, including M_2_-muscarinic, GABA_B_, A_1_-adenosine, α_2_-adrenergic, D_2_-dopamine receptors, as well as 5-HT_1A_-serotonin, somatostatin, galanin, sphingosine-1-phosphate, opioid, and cannabinoid receptors (CB_1_, CB_2_). Through these GPCRs, GIRK channels mediate the effects of neurotransmitters, neuromodulators, and (neuro)hormones, playing a crucial role in maintaining neuronal homeostasis and regulating synaptic plasticity processes essential for learning, memory, hormone and pain signaling [1–5].

The precise regulation of GIRK channels by PIP_2_ and GPCRs is needed for maintaining proper cellular function [3, 6, 7]. The binding of PIP_2_ at the interface between transmembrane domain (TMD) and cytoplasmic domain (CTD) of GIRK channel is necessary for its function. The search for the PIP_2_-binding site identified a set of basic residues relatively conserved between all inwardly rectifying potassium channels. In GIRK2, these residues include: Lys64, Lys90, Trp91, Arg92, Lys194, Lys199 and Lys200 [8–13]. They are located close to cholesterol-binding site, with some residues binding to both PIP_2_ and cholesterol [14, 15].

Recently, GIRK channels have emerged as promising targets for therapeutic interventions, since dysregulation of their activity has been implicated in a range of pathological conditions related to nociception [16–18] and excitatory/inhibitory balance disruption in epilepsy [19–22], Alzheimer’s and Parkinson’s disease [23–28], addiction, mood disorders and attention deficit hyperactivity disorder [21, 29–34]. For over a decade, it has been established that various modulators of GIRK channels interact near residues involved in PIP_2_ binding, making these channels highly druggable target [35]. However, no drug was developed that would specifically target GIRK-PIP_2_ interaction. The aim of this work was to move one step closer towards targeting GIRK channels at PIP_2_-binding site (Graphical Abstract). High-throughput virtual screening (HTVS) of over one million compounds was performed using Glide (Maestro), followed by SP/XP docking with RMSD-based core constraints, MM-GBSA binding free energy calculations, and optimization of pharmacokinetic and physicochemical properties. Finally, molecular dynamics (MD) simulations were done for 200 ns to evaluate binding dynamics and stability of protein-ligand complexes.

## 2. METHODS

### 2.1. Ligand preparation

A wide variety of synthetic and naturally ocurring molecules from almost all areas of life (bacterial, fungal, plant, animal) was obtained from publicly available compound databases. This included: PubChem [36], ZINC15 [37], CHEMBL [38], ASINEX [39], Collection of Open Natural Products [40], Pitt Quantum repository [41], Indian Medicinal Plants, Phytochemistry And Therapeutics [42], Peruvian Natural Products Database [43], Natural Products Atlas 3 [44], NuBBE DB [45], Medicinal Fungi Secondary Metabolite And Therapeutics [46], Comprehensive Marine Natural Products Database [47], and Human Metabolome Database [48]. The compounds were prepared using the LigPrep application in Maestro 13.5. Tautomeric and protonation states at pH = 7.4 ± 2.0 were generated using Epik [49], followed by ligand desaltation. An OPLS4 force field [50] was used to minimize the compounds. No more than 32 stereoisomers were generated for each ligand, retaining specified chiralities and varying other stereocenters.

### 2.2. Protein preparation

Homotetrameric GIRK2 crystal structure was obtained from the RCSB Protein Data Bank (PDB ID: 4KFM). The model system was built in Maestro (Schrödinger, LLC), by adding the missing side chains of residues, including Ile55, Arg73, Glu127, Phe141, Lys165, Lys301, and Glu303 in GIRK2; Arg42 and Arg214 in the Gβ subunit; as well as Glu58 and Glu63–Phe67 in the Gγ subunit. Protein Preparation module [51] was used to (1) fill missing short loops using Prime, (2) generate het states by Epik at the physiological pH = 7.4 ± 2, (3) remove any water molecules beyond 5 Å of het groups and (4) add missing hydrogens and determine the protonated states of the ionizable residues at pH = 7.4 by PROPKA pK_a_ calculations [52]. Finally, the GIRK2 structure was optimized with the OPLS4 force field.

### 2.3. Molecular docking

The Glide application in Maestro [53, 54] was used to conduct docking simulations. Protein grid was generated using the Receptor Grid Generation module, restricting docking to a 32 Å box centered on the co-crystallized native ligands. Docking with flexible ligand sampling was performed in three precision steps. All compounds were initially screened using the HTVS protocol, which allowed for the rapid evaluation of over one million compounds from publicly available databases, generating at most 32 poses per ligand. Poses were excluded if the sum of their Coulomb and van der Waals interaction energies exceeded 0 kcal / mol. Post-docking minimization and Epik state penalties were applied to docking score, and compounds with scores < −4.00 (more negative values indicate stronger predicted binding affinity) were selected for further refinement. Redocking was then performed with Standard Precision (SP) and Extra Precision (XP), and the final output score was expressed as the XP Glide Score (XP GScore) in kcal / mol.

In order to warrant the accuracy and relevance of the results, the docking protocol was guided to prioritize poses that are more likely to interact with key residues at the phosphoinositide-binding site. This was achieved by restricting docking to reference position with RMSD-based core constraint (< 2 Å of tolerance), requiring the core of each docked ligand to remain structurally similar to a reference native ligand.

### 2.4. Binding free energy calculation

To further assess the stability and strength of protein-ligand interactions, we performed binding free energy calculations using the Molecular Mechanics Generalized Born Surface Area solvation (MM-GBSA) method with all flexible residues (Prime). The OPLS4 force field and the variable-dielectric generalized Born model (VSGB) refinement solvation model were chosen to predict the binding free energy of complexes according to the following formula:

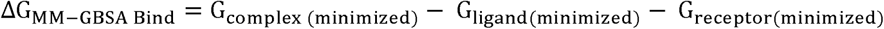

Ligands with binding free energy (higher affinity) lower than −100 kcal / mol were prioritized for further evaluation.

### 2.5. ADMET filtering and ligand optimization of known drugs / drug-like molecules

QikProp application in Maestro was used to predict pharmacokinetics, ADMET properties and druglikeness features of all drug-like molecules (excluding human metabolites). This included evaluating the number of property or descriptor values that fall outside the 95% range of similar values for known drugs, Lipinski rule of five (RO5) and Jorgensen’s Rule of three (RO3)), polar surface area (PSA), octanol/water partition coefficient (QPlogPo/w), CNS activity, blood-brain barrier (BBB) permeability (compounds with QPPMDCK < 25 were removed), gut-blood permeability (those with QPPCaco < 25 were excluded), and percent of human oral absorption. Potential toxicity and off-target propensity were examined by the number of metabolites and reactive functional groups, and the predicted IC_50_ value for blockage of HERG (QPlogHERG), cardiac rapidly activating delayed rectifier K^+^ channels that also bind PIP_2_. Compounds with > 8 metabolites, > 2 reactive functional groups and QPlogHERG values < −5 were deprioritized, as potential concern of cardiotoxicity. Finally, the Physics-Based Membrane Permeability module in Maestro was used to estimate the energy required for a compound to permeate the membrane (Membrane ΔG_Insert_).

The optimization of compounds with undesired properties was performed at each step (2.3 – 2.5). The properties optimized in this way included: XP GScore, binding free energy (ΔG_MM-GBSA Bind_), and ADMET properties, particularly oral bioavailability, CNS activity, QPlogHERG and gut-blood, BBB and membrane permeabilities. Additional compounds were obtained through: (1) similarity searches (Tanimoto > 80 %) and (2) bioisostere replacement with no immutable regions. Thus obtained compounds were then analyzed using the same protocol (2.3 – 2.5), and the most promising ligands were added to the set of compounds to be evaluated by molecular dynamics.

### 2.6. Molecular dynamics

Molecular dynamics (MD) simulations were conducted using GROMACS v2020.1 to investigate protein-ligand stability and ligand-induced dynamics [55]. MD simulations lasting 200 ns were done for apoprotein form of GIRK2 (without ligand), GIRK2-PIP_2_ docked complex, and best-scored protein-ligand complexes. Each system was embedded in a POPC (16:0, 18:1 PC) lipid bilayer (CHARMM-GUI membrane builder) and solvated with TIP3P water and 150 mM KCl. The CHARMM36 force field [56] was applied, with five KL ions in the conduction pathway and four NaL ions in sodium-binding sites (as in the crystal structure). Systems were equilibrated for 20 ns with Cα restraints (1,000 kJ/mol/nm²), followed by 200 ns unrestrained production runs.

Simulations used a 2-fs time step with periodic boundary conditions in all directions. Temperature and semi-isotropic pressure (compressibility = 4.5 × 10LL barL¹) were maintained using the velocity-rescale thermostat and Parrinello-Rahman barostat. Electrostatic interactions were calculated via the particle mesh Ewald (PME) method with a 0.12-nm Fourier spacing, and LINCS constraints were applied to all bonds. MD trajectories were analyzed for various parameters, including root mean square deviation (RMSD), root mean square fluctuation (RMSF, the time average of the RMSD), radius of gyration (Rg), and the number of intermolecular hydrogen bonds (H-bonds) between GIRK2 and each ligand.

## 3. RESULTS

### 3.1. Molecular docking

Redocking the core structure of the native ligand (PIP_2_ without acyl chains; PDB ID: PIO) to the GIRK2 channel yielded a binding free energy (ΔG_MM-GBSA Bind_) of −144.14 kcal / mol and a root-mean-square deviation (RMSD) of 1.05 Å (Fig. 1B, E). These results demonstrate a strong binding affinity and high accuracy in reproducing the native ligand binding pose (Fig. 1A). In comparison, molecular docking of the full native ligand (PIP_2_ with complete acyl chains, Fig. 1C, F) produced even stronger affinity (lower binding free energy of −176.85 kcal / mol) and shorter hydrogen bonds between the ligand and the receptor (< 2 Å, Fig. 1F), when compared to the core ligand (Fig. 1E). This suggests that the longer acyl chains of PIP_2_ may strengthen interactions with GIRK2, which is consistent with previous experimental observations [57–59]. In both docked poses, the key amino acids involved in PIP_2_ recognition were engaged: Lys64 (N-terminus), Lys90, Trp91, Arg92 (post-slide helix), Lys194 (cytosolic end of transmembrane helix), Lys199 and Lys200 (parts of the tether helix or C-linker) [8, 9, 11, 12]. Moreover, the docking protocol additionally recruited Gln197 from the β-loop of the GIRK2 channel, an interaction not present in the crystallographic structure (PDB ID: 4KFM). As a part of the C-linker (residues: 197-203) [14], Gln197 is involved in GIRK channel gating through the allosteric modulation by both PIP_2_ and cholesterol [15, 60].

**Figure 1.**
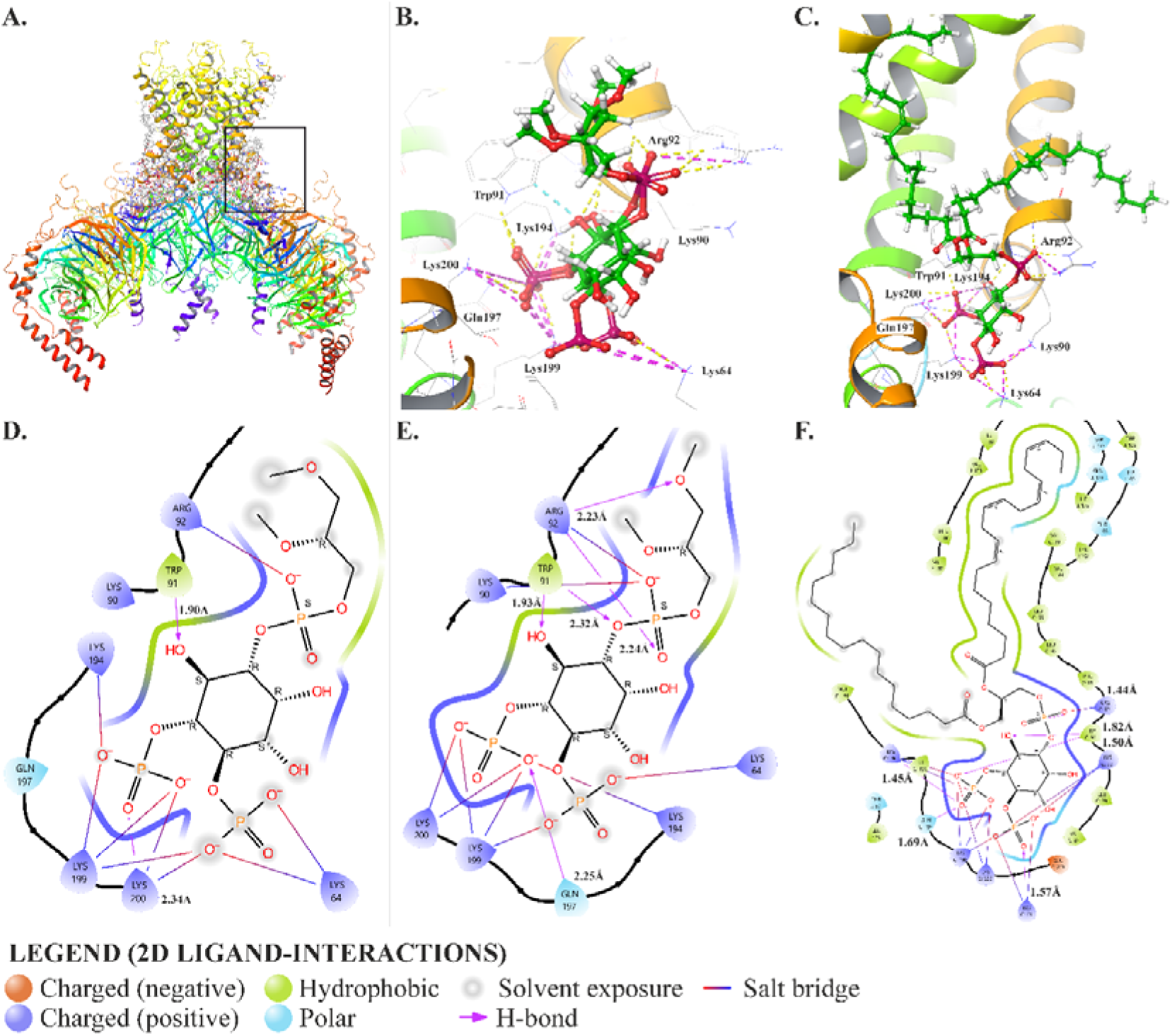
Validation of molecular docking protocol upon native ligand redocking against the GIRK2 homotetramer (**A**, PDB ID: 4KFM). **B.** Superposition of the native ligand pose present in the crystallographic structure **(D**, PDB ID: PIO, core PIP_2_ structure without acyl chains) and the obtained docked pose of the same ligand (**E**, ΔG_MM-GBSA Bind_ = −144.14 kcal / mol) revealed root-mean-square deviation of atomic positions (RMSD) of 1.05 Å, validating the docking protocol. **C**, **F.** Docked pose of full native ligand (PIP_2_ with complete acyl chains; 1-stearoyl-2-arachidonoyl-sn-glycero-3-phospho-(1′-myo-inositol-4′,5′-bisphosphate)) with the ΔG_MM-GBSA Bind_ = −176.85 kcal / mol.

The initial HTVS protocol included screening over 1,134,542 compounds for the GIRK-PIP_2_ binding site (Table S1), with 398,680 compounds further evaluated in XP mode (Table S2), followed by MM-GBSA calculations. Upon filtering, the best-scored compounds were classified into known drugs/drug-like molecules and human metabolites (Table 1, Fig. 2). The most promising drugs and drug-like molecules were then evaluated with MD simulations during 200 ns (Table S3).

**Figure 2.**
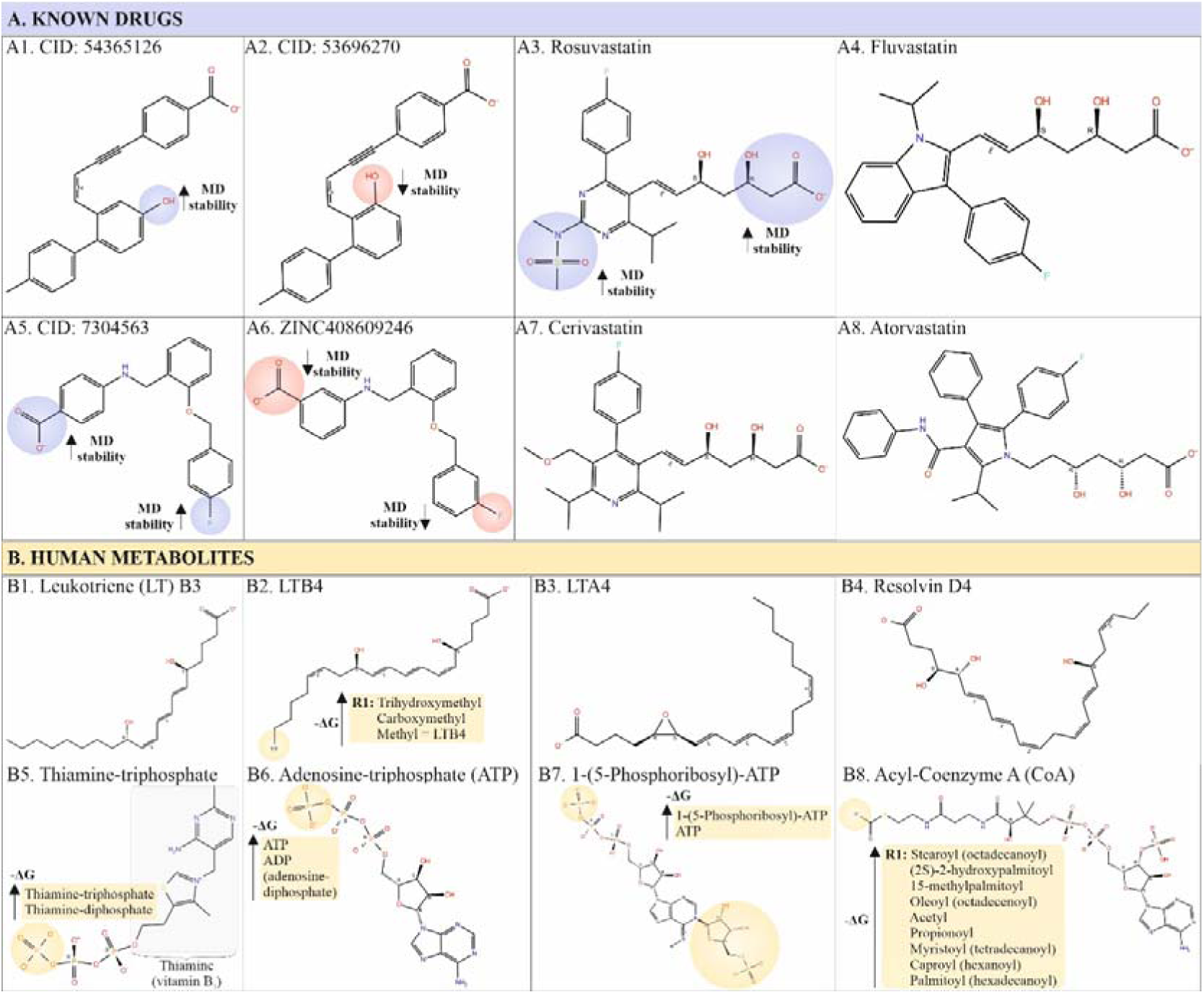
Chemical structures of the top 10 best-scored **A.** known drugs/drug-like compounds and **B.** human metabolites. The circles within the structures represent the relationships between ligand structure and **A.** ligand-GIRK2 complex stability during 200 ns of MD simulation for known drugs (blue/red), and **B.** post-docking binding free energy estimates for human metabolites (beige), with higher −ΔG (upward pointing arrows) indicating stronger binding affinity for the given order of substituents/structures.

**Table 1.** Docking results for GIRK2 channel (PDB ID: 4KFM) at the binding site of phosphatidylinositol-4,5-bisphosphate (PIP_2_) showing best-scored known drugs and drug-like molecules (A1-A6), and (B) human metabolites (B1-B6). Abbreviations: CoA – Coenzyme A; ΔG – Binding Free Energy (MM-GBSA); dock. – docking; DPA – docosapentaenoyl; OH – hydroxy; LT – leukotriene; XP GScore – Extra Precision Glide Score.

| **A. Known drugs / drug like molecules** |  |  |  |  |  |
| --- | --- | --- | --- | --- | --- |
| **Ligand (Chemical ID, CID)** |  | **dock. score** | **XP GScore** | **$\Delta G$** | **GIRK2 interacting residues** |
| A1 | 54365126 | -8.488 | -8.492 | -204.52 | Thr80, Lys90, Trp91, Arg92, Lys 200, Phe186 |
| A2 | 53696270 | -6.742 | -6.745 | -161.54 | Lys90, Arg92, Phe186 |
| A3 | Rosuvastatin | -5.545 | -5.546 | -166.93 | Lys90, Trp91, Arg92 |
| A4 | Fluvastatin | -5.477 | -5.478 | -125.48 | Lys90, Trp91, Arg92, Phe186 |
| A5 | 7304563 | -6.920 | -6.920 | -166.92 | Lys90, Trp91, Arg92 |
| A6 | ZINC408609246 | -6.669 | -6.669 | -153.62 | Trp91, Lys194, Lys 200 |
| A7 | Cerivastatin | -5.479 | -5.490 | -152.40 | Lys90, Trp91, Arg92 |
| A8 | Atorvastatin | -5.777 | -5.778 | -122.77 | Lys90, Trp91, Arg92, Phe186 |
| **B. Human metabolites** |  |  |  |  |  |
| B1 | Leukotriene B3 | -6.226 | -6.228 | -185.74 | Lys90, Arg92, Lys200, Thr80 |
| B2 | Leukotriene B4 | -7.339 | -7.340 | -168.55 | Arg92, Lys200, Thr80 |
| B3 | Leukotriene A4 | -6.195 | -6.195 | -159.21 | Arg92 |
| B4 | Resolvin D4 | -6.352 | -6.353 | -156.14 | Lys90, Arg92, Lys200, Thr80 |
| B5 | Thiamine-triphosphate (TTP) | -5.038 | -5.080 | -195.61 | Leu79, Lys90, Trp91, Arg92, Lys194, Lys200 |
| B6 | Adenosine-triphosphate (ATP) | -5.874 | -5.907 | -125.98 | Lys64, Lys90, Trp91, Arg92, Lys194 |
| B7 | 1-(5-phosphoribosyl)-ATP | -6.350 | -7.209 | -173.77 | Lys64, Lys90, Trp91, Arg92, Lys194, Gln197, Lys199, Lys200; Thr80 |
| B8 | Stearoyl-CoA | -8.707 | -10.474 | -206.25 | Lys64, Asp88, Lys90, Trp91, Arg92, Lys194, Lys199, Lys200, Asn231, Glu315 |

#### 3.1.1. Known drugs / drug-like molecules

Among known drugs, several statins and other similar molecules exhibited strong binding to the key PIP2-interacting residues. These compounds were selected for further validation via MD simulation analysis. The most promising binding was found for A1 (CID: 54365126, 4-[4-[5-hydroxy-2-(4-methylphenyl)phenyl]but-3-en-1-ynyl]benzoic acid), followed by A5 (CID: 7304563, 4-[[2-[(4-fluorophenyl)methoxy]phenyl]methylamino]benzoate) and A3 (Rosuvastatin). Strong binding was also found for similar compounds, including Cerivastatin, Fluvastatin and Atorvastatin (Table 1A, Fig. 3).

**Figure 3.**
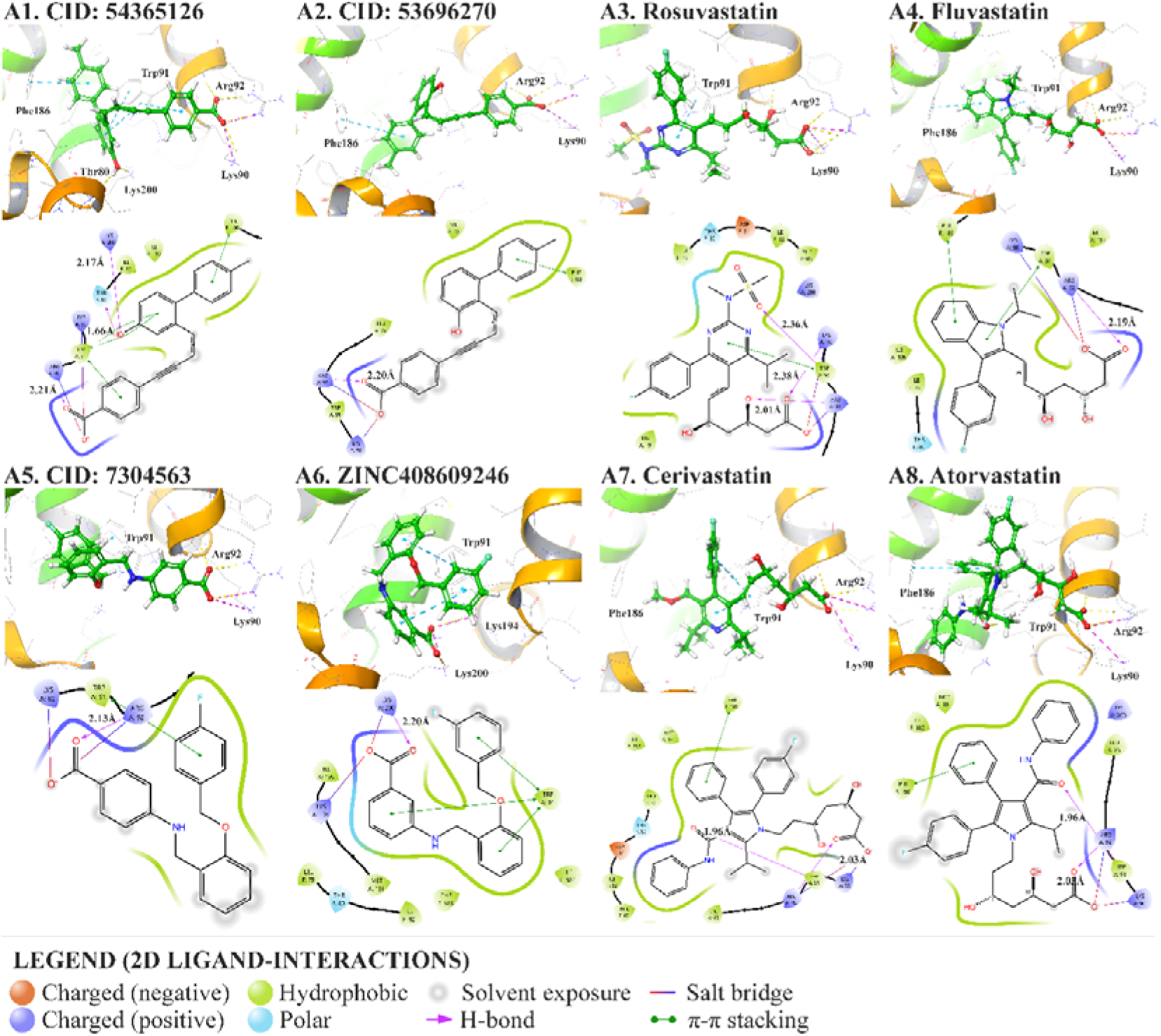
3D and 2D GIRK2-ligand interactions at PIP_2_-binding site (4 Å cut-off) for the 8 best-scored known drugs and drug-like molecules. The distances of pure H-bonds (not including salt bridges, labeled as purple arrows) are indicated in Å.

For all known drug-like molecules (A1-A6), favorable interactions were observed with key residues at the PIP_2_-binding site of GIRK2 (Table 1A, Fig. 3). These interactions included strong H-bonds and salt bridges (< 2.4 Å) between positively charged Lys90 and Arg92 and oxygen atoms of ligands, and frequent π-π stacking with Trp91 (6/8 ligands). Nearly all statins engaged in π-π stacking interactions with Trp91 at the PIP_2_ site, while Atorvastatin specifically formed π-π stacking interactions with Phe186, residue located at the cholesterol-binding site B. This site is situated in very close proximity to the PIP_2_-binding site and encompasses both Trp91 and Phe186 [14, 61]. Fluvastatin and the top-scored compound (A1) had π-π stacking with both Trp91 and Phe186. Additionally, A1 formed strong H-bonds with Lys200 (2.17 Å), another key residue involved in PIP_2_ binding, as well as with Thr80 located in the transmembrane region TM1 (1.66 Å). Thr80 was recently reported to be involved in direct inhibition of GIRK channels by sigma-1 receptor antagonist BD1047 [62].

As listed in Table 2A, the top six known drugs exhibit significant variability in BBB permeability (QPPMDCK). Poor membrane and BBB permeability is especially notable for Rosuvastatin (QPPMDCK below 25 nm / s and Membrane ΔG_Insert_ of 20.65 kcal / mol). A5 showed excellent human oral absorption and the most favorable BBB permeability, however, far from ideal, with QPPMDCK of 214.39 nm / s.

**Table 2.**
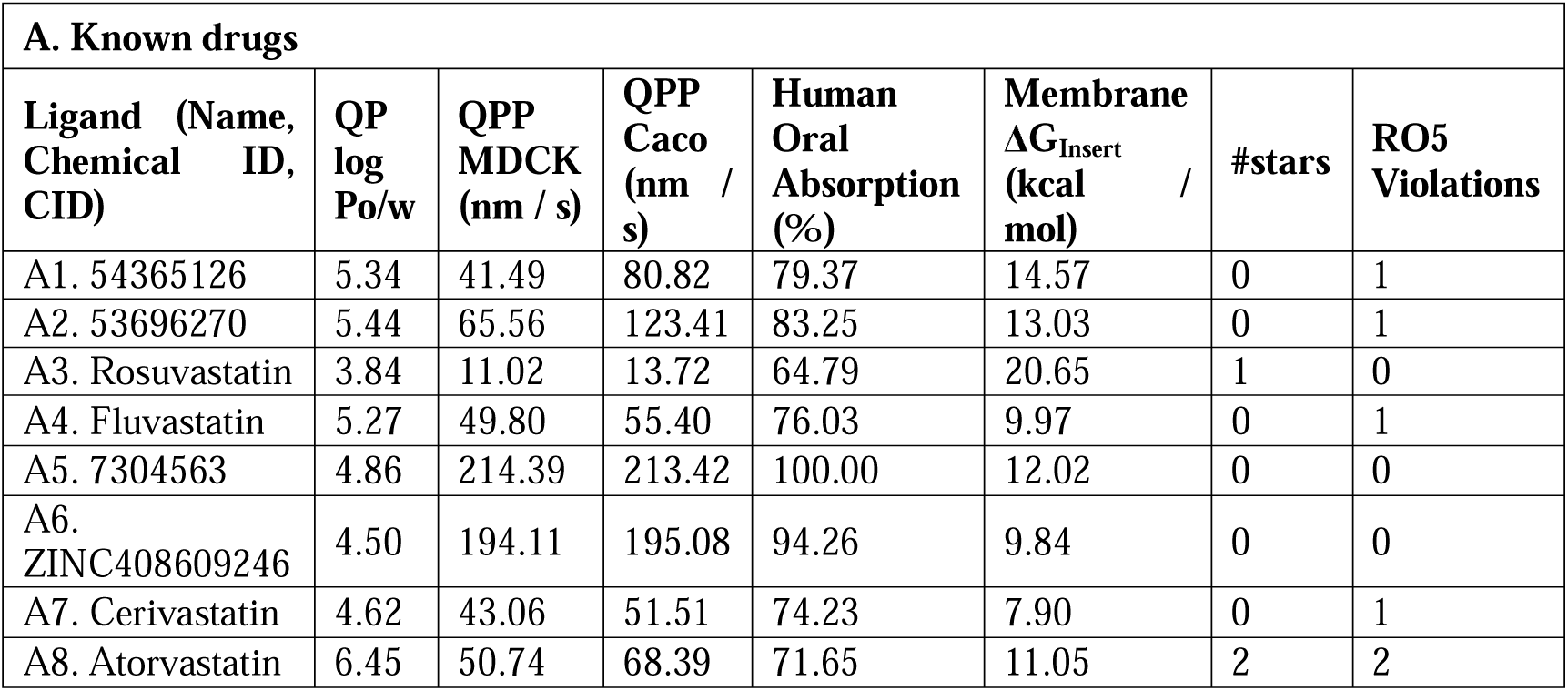
Important pharmacokinetic and physicochemical properties of the 8 best-scored ligands. Abbreviations: #stars - number of property or descriptor values falling outside the 95% range of similar values for known drugs; Membrane ΔG_Insert_ - the energy required for a compound to penetrate the cell membrane, with lower values indicating greater penetrability; QPlog Po/w - predicted octanol/water partition coefficient, larger values indicate higher lipophilicity; QPP Caco - predicted apparent Caco-2 cell permeability, for non-active transport. Caco-2 cells are a model for the gut-blood barrier (QPPCaco < 25 poor permeability, > 500 excellent permeability); QPP MDCK - predicted apparent Madin-Darby canine kidney cells (MDCK) cell permeability, for non-active transport. MDCK cells are considered to be a good mimic for the blood-brain barrier (BBB). Similarly to QPPCaco, values of QPPMDCK below 25 suggest poor BBB permeability, and those above 500 excellent permeability. RO5 – Lipinski’s Rule of five.

| **A. Known drugs** |  |  |  |  |  |  |  |
| --- | --- | --- | --- | --- | --- | --- | --- |
| **Ligand (Name, Chemical ID, CID)** | **QP log Po/w** | **QPP MDCK (nm / s)** | **QPP Caco (nm / s)** | **Human Oral Absorption (%)** | **Membrane $\Delta G_{\text{Insert}}$ (kcal / mol)** | **#stars** | **RO5 Violations** |
| A1. 54365126 | 5.34 | 41.49 | 80.82 | 79.37 | 14.57 | 0 | 1 |
| A2. 53696270 | 5.44 | 65.56 | 123.41 | 83.25 | 13.03 | 0 | 1 |
| A3. Rosuvastatin | 3.84 | 11.02 | 13.72 | 64.79 | 20.65 | 1 | 0 |
| A4. Fluvastatin | 5.27 | 49.80 | 55.40 | 76.03 | 9.97 | 0 | 1 |
| A5. 7304563 | 4.86 | 214.39 | 213.42 | 100.00 | 12.02 | 0 | 0 |
| A6. ZINC408609246 | 4.50 | 194.11 | 195.08 | 94.26 | 9.84 | 0 | 0 |
| A7. Cerivastatin | 4.62 | 43.06 | 51.51 | 74.23 | 7.90 | 0 | 1 |
| A8. Atorvastatin | 6.45 | 50.74 | 68.39 | 71.65 | 11.05 | 2 | 2 |

Among the known drugs with strong affinity for GIRK2 that were not included in Table 2A due to their low selectivity and undesirable ADMET and physicochemical properties, the most notable examples include: (1) ivermectin (docking score −4.130, ΔG −179.66 kcal / mol), already reported to activate GIRK2 at PIP_2_-site, with greater efficacy in activating GIRK2 than GIRK4 [63], (2) novel compound PG-21 (docking score −4.114, ΔG −156.54 kcal / mol); two cephalosporin antibiotics, namely (3) cefonicid (docking score −5.681, ΔG −173.66 kcal / mol), (4) carbacefamandole (docking score −5.771, ΔG −144.69 kcal / mol)) and (5) fungal metabolite zaragozic acid B (docking score −5.294, ΔG −133.66 kcal / mol.

PG-21 (Fig. S1) is a specifically designed PROTAC (proteolysis-targeting chimera) molecule, aimed to target and induce degradation of glycogen synthase kinase-3β (GSK-3β) in Alzheimer’s disease [64]. This compound showed strong van der Waals and electrostatic interactions with GIRK2, establishing two H-bonds with Trp91 and Arg92 at PIP_2_-binding site, π-cation interactions with Arg92, as well as π-stacking with cholesterol-binding Phe186.

However, it exhibited some unfavorable properties, including low BBB permeability (QPPMDCK only 2.96 nm / s) and potential cardiotoxicity (QPlogHERG of −7.9, with concerns for values below −5).

Cephalosporins (Fig. S2 and S3) were excluded due to their extremely low oral bioavailability and BBB permeability. Yet, it should be noted that these antibiotics require further investigation, as they have well-known effects on the central nervous system, being able to enter from blood into neurons in case of BBB disruption, either due to inflammation, vascular impairment or neurodegeneration [65–69].

Finally, zaragozic acid B (squalestatin, CID: 9940176), was excluded due to its poor pharmacokinetic properties, including high polarity and susceptibility to hydrolysis under acidic and basic conditions. This compound is a well-characterized inhibitor of mammalian squalene synthase, an enzyme involved in cholesterol biosynthesis [70, 71]. A closer look at the protein-ligand interaction analysis (Fig. S25) reveals that zaragozic acid B may mimic the binding characteristics of PIP_2_. Specifically, its 4,8-dioxabicyclo[3.2.1]octane core closely resembles the inositol ring of PIP_2_, both in terms of ligand geometry and by engaging with the same set of critical residues within the pocket (Fig. S26).

#### 3.1.2. Human metabolites

Among the human metabolites, many lipophilic signalling molecules, such as numerous acyl-CoA, polyphosphate compounds, leukotrienes LTA4, LTB3, LTB4, and their analogues showed high affinity for GIRK2 (Fig. 2B, 4 and S6-S23).

**Figure 4.**
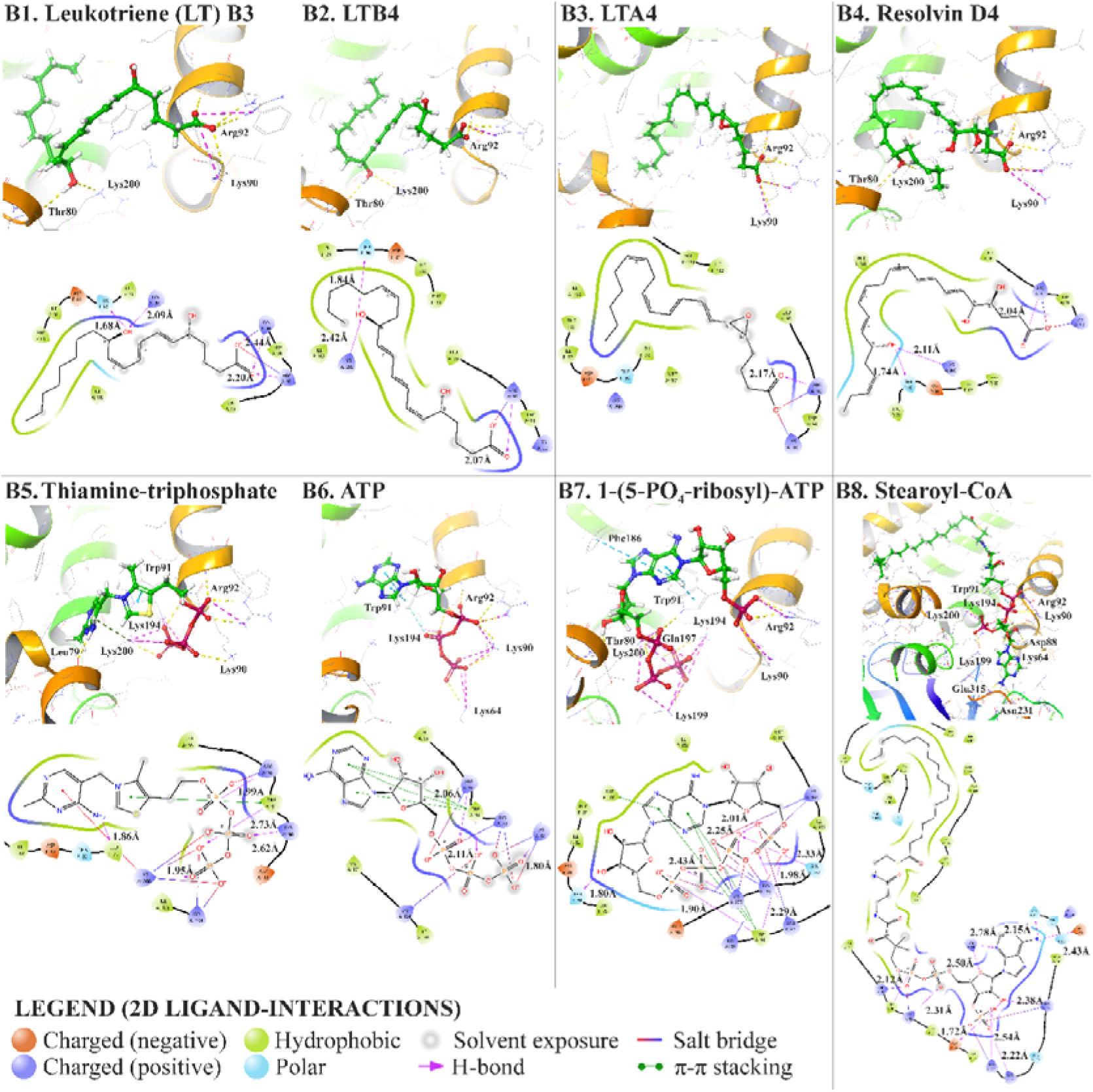
3D and 2D GIRK2-ligand interactions at PIP_2_-binding site (4 Å cut-off) for 8 best-scored human metabolites. The distances of the pure H-bonds (not including salt bridges, labeled as purple arrows) are indicated in Å. Abbreviations: ATP – adenosine-triphosphate, CoA – coenzyme A; LT – leukotriene, PO_4_ - phospho.

Stearoyl-CoA (B8) showed the strongest binding to GIRK2 among all acyl-CoAs (Table 1, Fig. 2B, Fig. S10-S22) and other metabolites analyzed in this work. The analysis of GIRK2-acyl-CoA interactions showed that the adenosine and phosphate moieties of CoA interacted closely (< 3 Å) with the key residues at the PIP_2_-binding site (Table 1B, Fig. 4): Lys64, Lys90, Trp91, Arg92, Lys199, Lys194 and Lys200. Furthermore, H-bonds were established between ribose moiety of stearoyl-CoA and Asp88, and between adenine nitrogen atoms and Asn231 (involved in Na^+^ coordination with GIRK [13]) and Glu315. Only hydrophobic interactions were found between GIRK2 and pantetheine moiety of CoA, with no H-bonds or salt bridges.

Several sterically similar polyphosphates were identified to bind to GIRK2 akin to the negatively charged phosphate groups of PIP_2_. The best-scored polyphosphates established complex interactions with almost all key residues at PIP_2_-binding site, and additionally recruited nearby Leu79, Thr80, and Phe186 (Fig. 4). These include adenosine-diphosphate (ADP), adenosine-triphosphate (ATP, B6 in Fig. 4), N1-(5-phospho-D-ribosyl)-ATP (B7 in Fig. 4), thiamine-diphosphate (also called thiamine pyrophosphate, TPP, the active (coenzyme) form of vitamin B1; Fig. S9) and thiamine-triphosphate (TTP, B5 in Fig. 4). Phosphorylation of ADP and TPP, as well as the addition of a 5-phosphoribosyl group to ATP, enhanced binding free energy estimates, whereas their monophosphates and dephosphorylated forms (adenine and thiamine or vitamin B_1_) showed no favorable binding within the 2 Å core-constraints. Likewise, adenosine 3’,5’-cyclic monophosphate (cAMP) exhibited very weak binding at the PIP_2_-site (−55.34 kcal / mol, Fig. S24).

Leukotrienes (LTs) and resolvins D series (resolution phase interaction products) exhibited a consistent binding profile with GIRK2. Almost all LTs and their derivatives established H-bonds and salt bridges with Lys90, Arg92, and Lys200 at the PIP_2_-binding site, and some engaged with Trp91 and Lys194 (Fig. 4). Four compounds additionally formed H-bonds with Phe83 and Ile195 near PIP_2_-binding site: resolvin E2 (Fig. S4), 12-oxo resolvin E1 (Fig. S5), 20-carboxy-LTB4 (Fig. S6) and 20-tri-OH-LTB4 (Fig. S7). These amino-acids are involved in GIRK channel activation by cholesterol (Phe83) [15] and ivermectin (Ile195 in the C-linker) [63]. Also 20-tri-OH-LTB4 formed H-bond with Ile182, another residue involved in cholesterol-GIRK interaction [15]. In addition, resolvin D4, LTB3, LTB4 (Fig. 4) and three LTB4 derivatives (12-oxo-20-tri-OH-LTB4, 20-tri-OH-LTB4 and 6-trans-LTB4; Fig. S23, S7, and S8, respectively) established one or two strong H-bonds (< 2.5 Å) with Thr80, residue involved in GIRK channel inhibition by the sigma-1 receptor antagonist BD1047 [62].

### 3.2. Molecular dynamics

Among the 32 GIRK2-drugs complexes subjected to molecular dynamics (MD) simulations (Table S3), three compounds exhibited stable binding with GIRK2 at PIP_2_ binding site. This stability was quantitatively assessed through low root-mean-square deviation (RMSD) values of the ligands and persistent H-bonds throughout the simulation trajectory (Fig. 5). The identified compounds were A1 (CID: 54365126, Fig. 5, depicted in blue), A3 (Rosuvastatin, red) and A5 (CID: 7304563, green). Among these, A5 showed the most favorable binding profile. No stable interactions with GIRK2 were found for other similar compounds, such as Fluvastatin, Atorvastatin, Cerivastatin and other molecules obtained through similarity searches and bioisostere replacements (Table S3). Based on these findings, the correlation between the ligand structure and the stability of GIRK2-ligand complex was depicted in Fig. 2A (blue/red).

**Figure 5.**
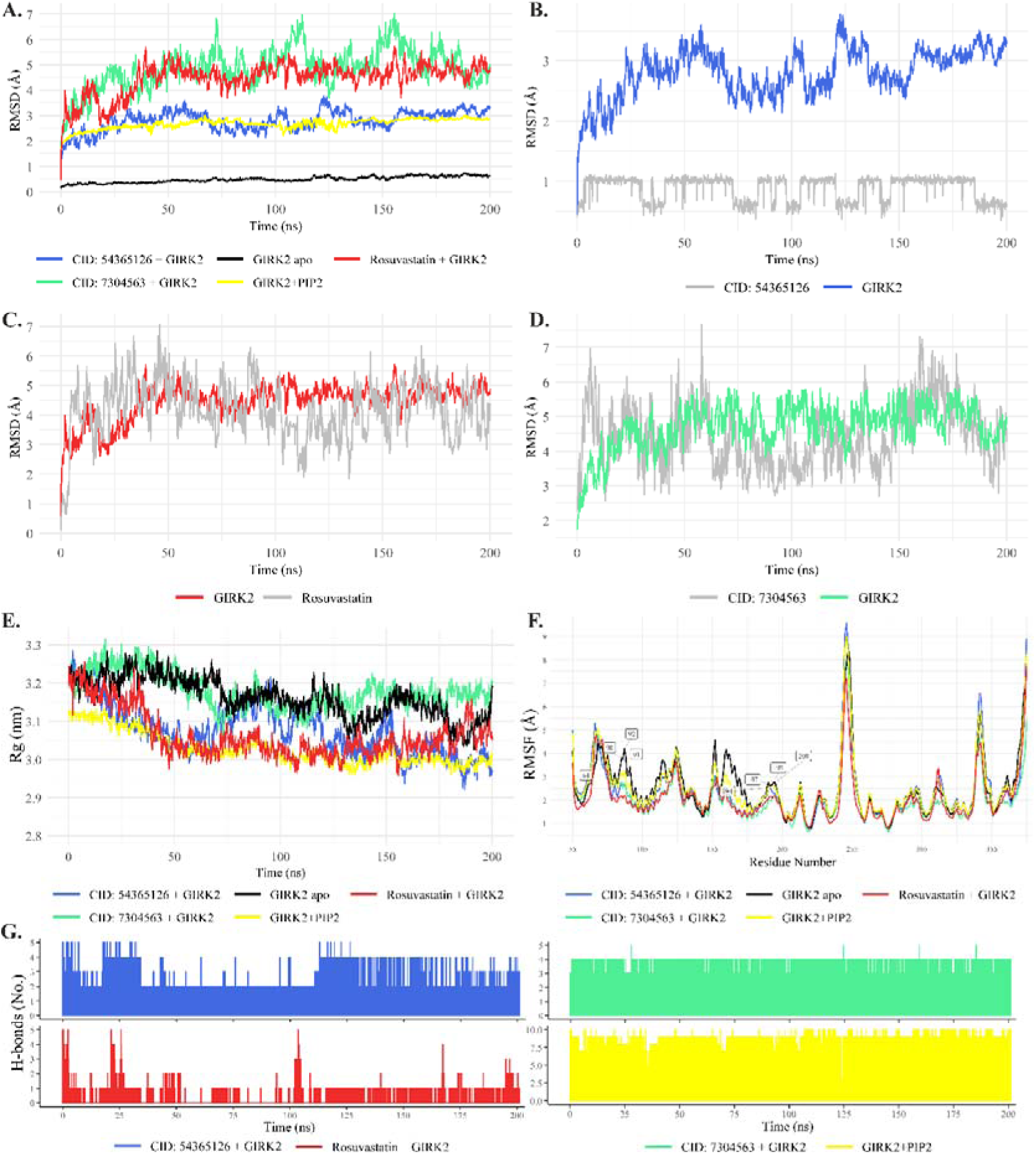
MD simulation analysis (200 ns). **A.** RMSD of apo GIRK2 (black), GIRK2–PIP2 (yellow), and GIRK2 bound to the top three drug complexes (see legend)**. B-D.** RMSD calculated separately for the GIRK2 protein backbone and heavy atoms of the top three drugs within the complex. **E.** Radius of gyration (Rg) and **F.** Root-mean-square-fluctuation (RMSF) of apo GIRK2, GIRK2 + PIP_2_ and top three drug-bound systems with labeled key PIP_2_-interacting residue numbers. **G.** Number of intermolecular hydrogen bonds (H-bonds) between GIRK2 and PIP_2_ (yellow) and top three drugs during 200 ns of simulation.

The apoprotein and docked GIRK-PIP_2_ complex exhibited high stability and minimal structural changes over 200 ns, with a backbone RMSD remaining well below 0.9 Å for apoprotein (Fig 5A, black) and below 2.9 Å for PIP_2_ (yellow). Upon ligand binding, structural perturbations were observed to varying degrees across the top three complexes (Fig. 5A, red, blue, green).

The complex with A1 (CID: 54365126, Fig. 5A blue) demonstrated the highest stability (RMSD ∼2.5 Å, similarly to PIP_2_), indicating tight binding with minor structural adjustments. Other two complexes exhibited greater conformational flexibility (RMSD ∼4–5 Å). When analyzed separately, the ligand-specific RMSD values further differentiated the binding modes of these ligands, with A1 (CID: 54365126, Fig. 5A blue) maintaining a near-rigid conformation (0.5–1 Å), and other two ligands having higher mobility that reflect ligand adjustments within the binding pocket.

The radius of gyration (Rg) analysis revealed a transient expansion of the apo protein (from 3.2 nm to 3.4 nm, Fig. 5E black) followed by re-compaction to its initial state. This suggests an initial relaxation phase followed by structural stabilization. The protein-ligand complexes displayed slightly different responses, with A5 (CID: 7304563, green) Rg values closely following those of apoprotein. On the other hand, binding of A1 (CID: 54365126, Fig. 5E blue) and A3 (Rosuvastatin, red) induced more significant compaction in the structure.

Root-mean-square fluctuations (RMSF) analysis further supported these observations. The apoprotein and all ligand-bound systems exhibited similar fluctuation profiles, with the highest flexibility observed in non-binding regions. In contrast, the key PIP_2_-interacting residues (Fig. 5F, square labels) exhibited reduced fluctuations in complex with top 3 ligands, suggesting that binding of these ligands may enhance stability of the key regions involved in PIP_2_ recognition.

Finally, the intermolecular H-bond analysis further elucidated the binding stability of each ligand. As can be seen on Fig. 5G, A5 (CID: 7304563, green) formed the most consistent interactions with GIRK2 at PIP_2_ binding site, maintaining 3–4 H-bonds throughout 200 ns. These findings align with the low RMSD and compact structure of this protein-ligand complex. Less stable interactions were found for the other two ligands, as reflected in their intermittent H-bond patterns and higher conformational flexibility. While A1 (CID: 54365126, blue) maintained at least two H-bonds throughout the simulation, A3 (Rosuvastatin, red) displayed intermittent hydrogen bonding with occasional complete bond loss, particularly during the first 100 ns, before achieving partial stabilization in later stages. This transient binding behavior suggests weaker initial interactions that gradually optimized over time, likely due to ligand repositioning within the binding pocket.

## 4. DISCUSSION

By integrating structure- and ligand-based approaches to drug discovery, this work identifies potential ligands with favorable and stable binding at PIP_2_-binding site of GIRK2. We found GIRK2-related effects of some known drugs and endogenous compounds, most notably statins, cephalosporins, leukotrienes, resolvins, TPP, ATP, as well as various acyl-CoAs and their metabolites. Direct interaction of these compounds with GIRK2 has not been previously reported; however, structurally related molecules are known to modulate similar channels or other GIRK subunits, supporting further investigation.

The identified compounds exhibited binding modes similar to the native ligand, interacting with critical PIP_2_-binding residues. Some ligands further established favorable interactions with residues responsible for binding cholesterol [14, 15] and GIRK-acting drugs, such as ivermectin [63] and sigma-1 receptor antagonist BD1047 [62]. Given the role of these residues in physiological and pharmacological channel regulation, the identified compounds may function as competitive GIRK modulators, offering a novel way to regulate channel activity.

### 4.1. Known drugs

Statins are widely used inhibitors of 3-hydroxy-3-methylglutaryl-CoA reductase, a critical enzyme in cholesterol biosynthesis. In the field of pharmacology, these drugs are somewhat (in)famous for their off-target effects that can have myopathic and hepatic consequences, leading to musculoskeletal changes with still unexplained etiology [72] and statin-induced liver injury [73], respectively. However, some off-target effects of statins could be desirable, including epigenetic [74], anti-inflammatory, anti-oxidant and endothelium repairing properties [75]. These beneficial effects led to numerous strategies of repurposing statins in conditions such as Alzheimer’s [76], diabetes mellitus [77], inflammatory disorders [78, 79] and cancer [78].

Atorvastatin administration and cholesterol intake are formerly reported to regulate GIRK channels, however, this was attributed to the effects of cholesterol on GIRK [80]. Also, it was shown that the application of simvastatin and mevastatin inhibit similar voltage-gated potassium channels Kv1.3 in cancer cells in a concentration-dependent manner. Mechanism of the inhibition probably includes direct interaction of statins with the Kv1.3 channel and perturbation of lipid bilayer structure that stabilizes the inactive state of the channels [81]. The co-administration of statins and flavonoids has been demonstrated to effectively block Kv1.3 channels, causing apoptosis in cancer cells [82]. Recently, Pitavastatin and Lovastatin were reported to inhibit vascular L-type calcium channels (Cav1.2) in rabbit aortic strips, enhancing amlodipine-induced vasorelaxation [83]. In contrast, Rosuvastatin, Simvastatin, Pravastatin and Atorvastatin were showed to stimulate Cav1.2 channels, leading to cerebral artery vasoconstriction in rats [84].

Here, among 32 drug-GIRK2 complexes assessed via 200 ns molecular dynamics simulations, three ligands, A1 (CID: 54365126), A3 (Rosuvastatin), and A5 (CID: 7304563), exhibited stable binding at the PIP2 site, as evidenced by low ligand RMSD and persistent hydrogen bonding. A5 showed the most favorable binding profile, forming 3-4 consistent H-bonds and maintaining compact structural integrity. A1 demonstrated tight binding with minimal flexibility, while A3 (Rosuvastatin) showed transient engagement that stabilized over time. Structural analyses (RMSD, Rg, RMSF) revealed ligand-induced compaction and reduced fluctuations at PIP_2_-site, supporting the role of these ligands in stabilizing GIRK2 conformation and potentially modulating its function. However, the stability of these interactions is far from ideal (Fig 5G, red), and further sampling with longer MD analyses and experimental studies are needed to confirm these findings. Basic bioisostere replacements and similarity searches did not further improve the binding or stability of these ligands, suggesting that they already occupy an optimal chemical space for these interactions. In contrast, the established docked poses for Fluvastatin, Atorvastatin and Cerivastatin failed to form stable complexes with GIRK2 above 100 ns (Table S3).

### 4.2. Human metabolites

Pro-inflammatory leukotrienes (LTs), anti-inflammatory resolvins, anionic polyphosphates, long-chain acyl-CoAs and their derivatives emerged as potential GIRK2 modulators, requiring further validation.

Of all acyl-CoAs investigated here (Fig. 2B, Fig. S10-S22), stearoyl-CoA (octadecanoyl-CoA) showed the strongest binding to GIRK2 (−206.25 kcal / mol). This is consistent with the docking results of the full native ligand and may reflect hydrophobic complementarity between the stearoyl chain and the binding pocket of PIP_2_ (1-stearoyl-2-arachidonoyl-sn-glycero-3-phospho-(1′-myo-inositol-4′,5′-bisphosphate), Fig. 1C, F). In addition, stearoyl/arachidonoyl species make up 50–80% of phosphoinositides (sn-1/sn-2) in primary mammalian cells [85–88]). The unsaturated analogue of stearoyl-CoA, oleoyl-CoA ((9Z)-octadecenoyl-CoA, Fig. S13, −172,66 kcal / mol), is already reported to competitively and reversibly inhibit all inwardly-rectifying potassium channels, including GIRKs [89]. Additionally, long-chain acyl-CoAs have been proposed to directly influence the PIP_2_ localization within the membrane [90], and gating mechanisms of other potassium and calcium channels [90–93]. Taken together, these findings point to an acyl chain-dependent membrane polarization. Furthermore, stearoyl-CoA in our model interacted with Asn231, residue involved in Na^+^ coordination with GIRK2 [13], suggesting that potential GIRK inhibition may be mediated by blocking intracellular Na^+^-dependent GIRK activation.

Somewhat unsurprisingly, several structurally similar polyphosphates (ADP, ATP, 1-(5-phosphoribosyl)-ATP, TPP, and TTP) were detected to interact with GIRK2 in a manner resembling the binding of phosphate groups from PIP_2_. These findings, if further validated, would point to a potential mechanism by which polyphosphates and other phosphomimetic compounds may modulate channel activity. By mimicking the native phosphoinositide head group, such molecules could interact with key residues responsible for lipid recognition, thereby influencing channel gating or stabilization. In case of adenosine-phosphates, previous studies found that ATP can enhance GIRK channel activity by activating ATP-dependent lipid kinases that generate PIP_2_ [94, 95]. Yet, the direct effect of ATP and similar compounds on GIRK has not been reported so far, requiring further investigation. As for vitamin B_1_ (thiamine), it is believed to be directly involved in nerve stimulation in a nonLcoenzymatic way due to its interference with the structure and function of cellular membranes and its ability to regulate ion channels [96–98]. Here, both coenzyme forms of vitamin B1, thiamine diphosphate (TPP) and thiamine triphosphate (TTP), exhibited strong binding modes that closely resembled that of PIP_2_. The additional phosphate group greatly improved binding affinity from TPP to TTP (ΔΔG_MM-GBSA_ = −50.98 kcal / mol). On the other hand, unphosphorylated adenine/thiamine and their monophosphates did not form a favorable docking pose within 2 Å RMSD of the native PIP_2_ core. However, alternative binding modes outside this constraint cannot be ruled out.

LTs are lipid mediators of inflammation and chemotaxis implicated in various disorders, including inflammatory, cardiovascular and neurodegenerative diseases [99, 100]. Arachidonic acid (present in the PIP_2_ structure) and some of its derivatives have been previously found to stimulate atrial K_ACh_ channel, composed of GIRK1 and GIRK4 [35, 101–103], but not GIRK2. Furthermore, LTB4 is known to modulate calcium- and voltage-gated potassium channels [104]. On the other hand, resolvins D1 and E1 can affect sensory TRP channels [105] and excitatory NMDA receptors involved in the spinal cord plasticity processes involved in pain signaling [106–109]. In our work, various LTs and three resolvins showed strong affinity towards key residues at the GIRK2-PIP_2_-binding site. Some of these compounds also interacted with nearby residues involved in channel regulation by cholesterol (Phe83, Ile182) [14], ivermectin (Ile195) [63] and the sigma-1 receptor antagonist BD1047 (Thr80) [62].

The binding profiles of leukotrienes and resolvins revealed here suggest a mechanistic overlap between these immune mediators and existing GIRK modulators. The cocktail of immune signalling molecules disrupting potassium conductance in neurons during an inflammatory event could ultimately affect cell excitability and even survival. This would be particularly important for immune response in neurodegenerative disorders, such as Alzheimer’s disease and Parkinson’s disease [100, 110–116], and other neurological and psychiatric conditions where both GIRK and immune signaling are impaired [25, 117–122]. Hence, a deeper mechanistic understanding of how immune lipids interact with neuronal ion channels may open up alternative strategies in analgesia, neurodegeneration and neuroimmune disorders [109, 116, 123–127]. It should be added that the lipid mediators analyzed here exhibit varying (and in some cases unknown) metabolic stability. Their metabolism involves oxidative inactivation by cytochrome P450 enzymes, including CYP4F2 [128–130]. Therefore, some mediators are rapidly degraded (e.g., those with epoxide ring, such as LTA4 [131]), resulting in transient effects, while others may last longer.

### 3.4. Study limitations and future directions

Here, we identified several known drugs and human metabolites as potential GIRK2 modulators. However, due to computational constraints, we prioritized validating only the known drugs using MD simulations, leaving the assessment of endogenous compounds for future studies. Further computational and experimental approaches will be necessary to confirm the effects of the identified compounds. Longer MD simulations (> 200 ns) could provide deeper insights into ligand stability, conformational changes, and binding dynamics. Given the tetrameric nature of GIRK channels and structural variations among subunits (GIRK1-4), homology modeling across subunits and species may help identify subunit- and species-specific ligand preferences. *In vitro* experiments, such as electrophysiological recordings and ligand-binding assays, would be essential to confirm the functional effects of ligand interactions. Patch-clamp electrophysiology, for instance, could be used to measure ion flow through the channels in response to ligand binding, while radioligand or fluorescence-based binding assays could determine the affinity and specificity of these interactions. Finally, *in vivo* studies would be critical to understanding the physiological relevance of these interactions by assessing the potential of these ligands in a biological context.

## 5. CONCLUSION

Molecular docking with RMSD-based core constraints and MD simulation analyses were used to identify novel modulators of GIRK2 channels targeting the PIP_2_-binding site. Three known compounds emerged as promising candidates able to form stable complexes with GIRK2 during 200 ns of MD simulation: Rosuvastatin, CID: 54365126 and 7304563. In addition, two antibiotics (cefonicid and carbacefamandole) and numerous well-known signalling molecules were identified as potential GIRK2 modulators, requiring further computational and experimental evaluation. These include leukotrienes, resolvins, acyl-CoAs, and polyphosphates. The binding patterns of acyl-CoAs and polyphosphates suggest a dependence on lipid and phosphate tails, respectively, mimicking the hydrophobic and anionic features of PIP_2_.

The ligands reported here engaged in favorable interactions with residues essential for binding physiological GIRK modulators, such as PIP_2_, cholesterol, and oleoyl-CoA, as well as some GIRK-acting drugs. This suggests significant potential for the identified compounds to modulate GIRK2 channel function by competing with endogenous modulators and pharmacological agents. Such modulation could have implications for regulating neuronal excitability and synaptic transmission in conditions where GIRK channels play a critical role, such as Alzheimer’s disease, addiction, pain signaling and mood disorders. Future research should validate these findings and further investigate the broader pharmacological potential of these compounds.

## Supporting information

Supplementary_GIRK2-ligand complexes

## AUTHORS’ CONTRIBUTIONS

JDNL and LJD were responsible for the initial conceptualization, funding acquisition, supervision, and project administration; DJ was responsible for Data curation, Software development, Formal analysis, visualization and writing the original draft; JD, JDNL and LJD did the writing – review and editing. All authors read and approved the final manuscript.

## SUPPLEMENTARY MATERIALS

1. Supplementary Figures S1-S26
2. Supplementary Tables S1, S2, S3
3. Supplementary Data - GIRK2-ligand complexes

## DATA AVAILABILITY

All data supporting this analysis is available.

## REFERENCES

[1] D. Jeremic, I. Sanchez-Rodriguez, L. Jimenez-Diaz, J.D. Navarro-Lopez, Therapeutic potential of targeting G protein-gated inwardly rectifying potassium (GIRK) channels in the central nervous system, Pharmacol Ther, 223 (2021) 107808.

[2] R. Luján, E. Marron Fernandez de Velasco, C. Aguado, K. Wickman, New insights into the therapeutic potential of Girk channels, Trends Neurosci, 37 (2014) 20–29.

[3] C. Lüscher, P.A. Slesinger, Emerging roles for G protein-gated inwardly rectifying potassium (GIRK) channels in health and disease, Nat Rev Neurosci, 11 (2010) 301–315.

[4] R. Ochi, Y. Momose, K. Oyama, W.R. Giles, Sphingosine-1-phosphate effects on guinea pig atrial myocytes: Alterations in action potentials and K+ currents, Cardiovasc Res, 70 (2006) 88–96.

[5] S. Constantin, S. Wray, Galanin Activates G Protein Gated Inwardly Rectifying Potassium Channels and Suppresses Kisspeptin-10 Activation of GnRH Neurons, Endocrinology, 157 (2016) 3197–3212.

[6] H. Hibino, A. Inanobe, K. Furutani, S. Murakami, I. Findlay, Y. Kurachi, Inwardly rectifying potassium channels: their structure, function, and physiological roles, Physiol Rev, 90 (2010) 291–366.

[7] H. Luo, E. Marron Fernandez de Velasco, K. Wickman, Neuronal G protein-gated K(+) channels, Am J Physiol Cell Physiol, 323 (2022) C439–c460.

[8] T. Rohács, J. Chen, G.D. Prestwich, D.E. Logothetis, Distinct specificities of inwardly rectifying K(+) channels for phosphoinositides, J Biol Chem, 274 (1999) 36065–36072.

[9] S. Pegan, C. Arrabit, W. Zhou, W. Kwiatkowski, A. Collins, P.A. Slesinger, S. Choe, Cytoplasmic domain structures of Kir2.1 and Kir3.1 show sites for modulating gating and rectification, Nat Neurosci, 8 (2005) 279–287.

[10] S.B. Hansen, X. Tao, R. MacKinnon, Structural basis of PIP2 activation of the classical inward rectifier K+ channel Kir2.2, Nature, 477 (2011) 495–498.

[11] C.M. Lopes, H. Zhang, T. Rohacs, T. Jin, J. Yang, D.E. Logothetis, Alterations in conserved Kir channel-PIP2 interactions underlie channelopathies, Neuron, 34 (2002) 933–944.

[12] H. Zhang, C. He, X. Yan, T. Mirshahi, D.E. Logothetis, Activation of inwardly rectifying K+ channels by distinct PtdIns(4,5)P2 interactions, Nat Cell Biol, 1 (1999) 183–188.

[13] M.R. Whorton, R. MacKinnon, Crystal structure of the mammalian GIRK2 K+ channel and gating regulation by G proteins, PIP2, and sodium, Cell, 147 (2011) 199–208.

[14] Y.K. Mathiharan, I.W. Glaaser, Y. Zhao, M.J. Robertson, G. Skiniotis, P.A. Slesinger, Structural insights into GIRK2 channel modulation by cholesterol and PIP(2), Cell Rep, 36 (2021) 109619.

[15] M. Cui, Y. Lu, X. Ma, D.E. Logothetis, Molecular mechanism of GIRK2 channel gating modulated by cholesteryl hemisuccinate, Front Physiol, 15 (2024) 1486362.

[16] D. Nockemann, M. Rouault, D. Labuz, P. Hublitz, K. McKnelly, F.C. Reis, C. Stein, P.A. Heppenstall, The K(+) channel GIRK2 is both necessary and sufficient for peripheral opioid-mediated analgesia, EMBO Mol Med, 5 (2013) 1263–1277.

[17] M.K. Chung, Y.S. Cho, Y.C. Bae, J. Lee, X. Zhang, J.Y. Ro, Peripheral G protein-coupled inwardly rectifying potassium channels are involved in δ-opioid receptor-mediated anti-hyperalgesia in rat masseter muscle, Eur J Pain, 18 (2014) 29–38.

[18] M. Kimura, H. Shiokawa, Y. Karashima, M. Sumie, S. Hoka, K. Yamaura, Antinociceptive effect of selective G protein-gated inwardly rectifying K+ channel agonist ML297 in the rat spinal cord, PLoS One, 15 (2020) e0239094.

[19] R.A. Rifkin, X. Wu, B. Pereira, B.J. Gill, E.M. Merricks, A.J. Michalak, A.R. Goldberg, N. Humala, A. Dovas, G. Rai, G.M. McKhann, P.A. Slesinger, P. Canoll, C. Schevon, A selective small-molecule agonist of G protein-gated inwardly-rectifying potassium channels reduces epileptiform activity in mouse models of tumor-associated and provoked seizures, Neuropharmacology, 265 (2025) 110259.

[20] Y. Zhao, P.M. Ung, G. Zahoránszky-Kőhalmi, A.V. Zakharov, N.J. Martinez, A. Simeonov, I.W. Glaaser, G. Rai, A. Schlessinger, J.J. Marugan, P.A. Slesinger, Identification of a G-Protein-Independent Activator of GIRK Channels, Cell Rep, 31 (2020) 107770.

[21] Y. Zhao, I. Gameiro-Ros, I.W. Glaaser, P.A. Slesinger, Advances in Targeting GIRK Channels in Disease, Trends Pharmacol Sci, 42 (2021) 203–215.

[22] Y. Huang, Y. Zhang, S. Kong, K. Zang, S. Jiang, L. Wan, L. Chen, G. Wang, M. Jiang, X. Wang, J. Hu, Y. Wang, GIRK1-mediated inwardly rectifying potassium current suppresses the epileptiform burst activities and the potential antiepileptic effect of ML297, Biomed Pharmacother, 101 (2018) 362–370.

[23] I. Sánchez-Rodríguez, S. Djebari, S. Temprano-Carazo, D. Vega-Avelaira, R. Jiménez-Herrera, G. Iborra-Lázaro, J. Yajeya, L. Jiménez-Díaz, J.D. Navarro-López, Hippocampal long-term synaptic depression and memory deficits induced in early amyloidopathy are prevented by enhancing G-protein-gated inwardly rectifying potassium channel activity, J Neurochem, 153 (2020) 362–376.

[24] S. Djebari, G. Iborra-Lázaro, S. Temprano-Carazo, I. Sánchez-Rodríguez, M.O. Nava-Mesa, A. Múnera, A. Gruart, J.M. Delgado-García, L. Jiménez-Díaz, J.D. Navarro-López, G-Protein-Gated Inwardly Rectifying Potassium (Kir3/GIRK) Channels Govern Synaptic Plasticity That Supports Hippocampal-Dependent Cognitive Functions in Male Mice, J Neurosci, 41 (2021) 7086–7102.

[25] A. Martín-Belmonte, C. Aguado, R. Alfaro-Ruiz, A.E. Moreno-Martínez, L. de la Ossa, E. Aso, L. Gómez-Acero, R. Shigemoto, Y. Fukazawa, F. Ciruela, R. Luján, Nanoscale alterations in GABA(B) receptors and GIRK channel organization on the hippocampus of APP/PS1 mice, Alzheimers Res Ther, 14 (2022) 136.

[26] O.J. Lieberman, F. Bartolini, M.C. Miniaci, GIRK channels in Alzheimer’s disease, Aging (Albany NY), 12 (2020) 18793–18794.

[27] S.P. Choudhury, S. Bano, S. Sen, K. Suchal, S. Kumar, F. Nikolajeff, S.K. Dey, V. Sharma, Altered neural cell junctions and ion-channels leading to disrupted neuron communication in Parkinson’s disease, NPJ Parkinsons Dis, 8 (2022) 66.

[28] A. Contreras, S. Djebari, S. Temprano-Carazo, A. Múnera, A. Gruart, J.M. Delgado-Garcia, L. Jiménez-Díaz, J.D. Navarro-López, Impairments in hippocampal oscillations accompany the loss of LTP induced by GIRK activity blockade, Neuropharmacology, 238 (2023) 109668.

[29] S. Dehbozorghi, S. Bagheri, K. Moradi, K. Shokraee, M.R. Mohammadi, S. Akhondzadeh, Efficacy and safety of tipepidine as adjunctive therapy in children with attention-deficit/hyperactivity disorder: Randomized, double-blind, placebo-controlled clinical trial, Psychiatry Clin Neurosci, 73 (2019) 690–696.

[30] H. Kotajima-Murakami, S. Ide, K. Ikeda, GIRK Channels as Candidate Targets for the Treatment of Substance Use Disorders, Biomedicines, 10 (2022).

[31] R.A. Rifkin, S.J. Moss, P.A. Slesinger, G Protein-Gated Potassium Channels: A Link to Drug Addiction, Trends Pharmacol Sci, 38 (2017) 378–392.

[32] J. Zhang, Y. Zhu, M. Zhang, J. Yan, Y. Zheng, L. Yao, Z. Li, Z. Shao, Y. Chen, Potassium channels in depression: emerging roles and potential targets, Cell Biosci, 14 (2024) 136.

[33] P. Imbrici, D.C. Camerino, D. Tricarico, Major channels involved in neuropsychiatric disorders and therapeutic perspectives, Front Genet, 4 (2013) 76.

[34] M. Okada, I. Kozaki, H. Honda, Antidepressive effect of an inward rectifier K+ channel blocker peptide, tertiapin-RQ, PLoS One, 15 (2020) e0233815.

[35] D.E. Logothetis, D. Lupyan, A. Rosenhouse-Dantsker, Diverse Kir modulators act in close proximity to residues implicated in phosphoinositide binding, J Physiol, 582 (2007) 953–965.

[36] S. Kim, J. Chen, T. Cheng, A. Gindulyte, J. He, S. He, Q. Li, B.A. Shoemaker, P.A. Thiessen, B. Yu, L. Zaslavsky, J. Zhang, E.E. Bolton, PubChem 2025 update, Nucleic Acids Res, 53 (2025) D1516–d1525.

[37] T. Sterling, J.J. Irwin, ZINC 15--Ligand Discovery for Everyone, J Chem Inf Model, 55 (2015) 2324–2337.

[38] D. Mendez, A. Gaulton, A.P. Bento, J. Chambers, M. De Veij, E. Félix, M.P. Magariños, J.F. Mosquera, P. Mutowo, M. Nowotka, M. Gordillo-Marañón, F. Hunter, L. Junco, G. Mugumbate, M. Rodriguez-Lopez, F. Atkinson, N. Bosc, C.J. Radoux, A. Segura-Cabrera, A. Hersey, A.R. Leach, ChEMBL: towards direct deposition of bioassay data, Nucleic Acids Res, 47 (2019) D930–d940.

[39] Asinex.com, Asinex Focused Libraries, Screening compounds, Pre-plated Sets. https://www.asinex.com/, 2025.

[40] M. Sorokina, P. Merseburger, K. Rajan, M.A. Yirik, C. Steinbeck, COCONUT online: Collection of Open Natural Products database, J Cheminform, 13 (2021) 2.

[41 ] R. Gupta, Pitt Quantum Repository, https://github.com/pittquantum, 2025.

[42] K. Mohanraj, B.S. Karthikeyan, R.P. Vivek-Ananth, R.P.B. Chand, S.R. Aparna, P. Mangalapandi, A. Samal, IMPPAT: A curated database of Indian Medicinal Plants, Phytochemistry And Therapeutics, Sci Rep, 8 (2018) 4329.

[43] H.L. Barazorda-Ccahuana, L.G. Ranilla, M.A. Candia-Puma, E.G. Cárcamo-Rodriguez, A.E. Centeno-Lopez, G. Davila-Del-Carpio, J.L. Medina-Franco, M.A. Chávez-Fumagalli, PeruNPDB: the Peruvian Natural Products Database for in silico drug screening, Sci Rep, 13 (2023) 7577.

[44] E.F. Poynton, J.A. van Santen, M. Pin, M.M. Contreras, E. McMann, J. Parra, B. Showalter, L. Zaroubi, K.R. Duncan, R.G. Linington, The Natural Products Atlas 3.0: extending the database of microbially derived natural products, Nucleic Acids Res, 53 (2025) D691–d699.

[45] A.C. Pilon, M. Valli, A.C. Dametto, M.E.F. Pinto, R.T. Freire, I. Castro-Gamboa, A.D. Andricopulo, V.S. Bolzani, NuBBE(DB): an updated database to uncover chemical and biological information from Brazilian biodiversity, Sci Rep, 7 (2017) 7215.

[46] R.P. Vivek-Ananth, A.K. Sahoo, K. Kumaravel, K. Mohanraj, A. Samal, MeFSAT: a curated natural product database specific to secondary metabolites of medicinal fungi, RSC Adv, 11 (2021) 2596–2607.

[47] C. Lyu, T. Chen, B. Qiang, N. Liu, H. Wang, L. Zhang, Z. Liu, CMNPD: a comprehensive marine natural products database towards facilitating drug discovery from the ocean, Nucleic Acids Res, 49 (2021) D509–d515.

[48] D.S. Wishart, A. Guo, E. Oler, F. Wang, A. Anjum, H. Peters, R. Dizon, Z. Sayeeda, S. Tian, B.L. Lee, M. Berjanskii, R. Mah, M. Yamamoto, J. Jovel, C. Torres-Calzada, M. Hiebert-Giesbrecht, V.W. Lui, D. Varshavi, D. Varshavi, D. Allen, D. Arndt, N. Khetarpal, A. Sivakumaran, K. Harford, S. Sanford, K. Yee, X. Cao, Z. Budinski, J. Liigand, L. Zhang, J. Zheng, R. Mandal, N. Karu, M. Dambrova, H.B. Schiöth, R. Greiner, V. Gautam, HMDB 5.0: the Human Metabolome Database for 2022, Nucleic Acids Res, 50 (2022) D622–d631.

[49] J.R. Greenwood, D. Calkins, A.P. Sullivan, J.C. Shelley, Towards the comprehensive, rapid, and accurate prediction of the favorable tautomeric states of drug-like molecules in aqueous solution, J Comput Aided Mol Des, 24 (2010) 591–604.

[50] C. Lu, C. Wu, D. Ghoreishi, W. Chen, L. Wang, W. Damm, G.A. Ross, M.K. Dahlgren, E. Russell, C.D. Von Bargen, R. Abel, R.A. Friesner, E.D. Harder, OPLS4: Improving Force Field Accuracy on Challenging Regimes of Chemical Space, J Chem Theory Comput, 17 (2021) 4291–4300.

[51] G.M. Sastry, M. Adzhigirey, T. Day, R. Annabhimoju, W. Sherman, Protein and ligand preparation: parameters, protocols, and influence on virtual screening enrichments, J Comput Aided Mol Des, 27 (2013) 221–234.

[52] D.C. Bas, D.M. Rogers, J.H. Jensen, Very fast prediction and rationalization of pKa values for protein-ligand complexes, Proteins, 73 (2008) 765–783.

[53] R.A. Friesner, J.L. Banks, R.B. Murphy, T.A. Halgren, J.J. Klicic, D.T. Mainz, M.P. Repasky, E.H. Knoll, M. Shelley, J.K. Perry, D.E. Shaw, P. Francis, P.S. Shenkin, Glide:L A New Approach for Rapid, Accurate Docking and Scoring. 1. Method and Assessment of Docking Accuracy, Journal of Medicinal Chemistry, 47 (2004) 1739–1749.

[54] M.P. Repasky, M. Shelley, R.A. Friesner, Flexible ligand docking with Glide, Curr Protoc Bioinformatics, Chapter 8 (2007) Unit 8.12.

[55] M.J. Abraham, T. Murtola, R. Schulz, S. Páll, J.C. Smith, B. Hess, E. Lindahl, GROMACS: High performance molecular simulations through multi-level parallelism from laptops to supercomputers, SoftwareX, 1 (2015) 19–25.

[56] R.B. Best, X. Zhu, J. Shim, P.E. Lopes, J. Mittal, M. Feig, A.D. MacKerell Jr, Optimization of the additive CHARMM all-atom protein force field targeting improved sampling of the backbone L, ψ and side-chain χ1 and χ2 dihedral angles, Journal of chemical theory and computation, 8 (2012) 3257–3273.

[57] J.W. Sohn, A. Lim, S.H. Lee, W.K. Ho, Decrease in PIP(2) channel interactions is the final common mechanism involved in PKC- and arachidonic acid-mediated inhibitions of GABA(B)-activated K+ current, J Physiol, 582 (2007) 1037–1046.

[58] E. Leal-Pinto, Y. Gómez-Llorente, S. Sundaram, Q.Y. Tang, T. Ivanova-Nikolova, R. Mahajan, L. Baki, Z. Zhang, J. Chavez, I. Ubarretxena-Belandia, D.E. Logothetis, Gating of a G protein-sensitive mammalian Kir3.1 prokaryotic Kir channel chimera in planar lipid bilayers, J Biol Chem, 285 (2010) 39790–39800.

[59] N. D’Avanzo, W.W. Cheng, D.A. Doyle, C.G. Nichols, Direct and specific activation of human inward rectifier K+ channels by membrane phosphatidylinositol 4,5-bisphosphate, J Biol Chem, 285 (2010) 37129–37132.

[60] A. Inanobe, Y. Kurachi, Membrane channels as integrators of G-protein-mediated signaling, Biochim Biophys Acta, 1838 (2014) 521–531.

[61] A.N. Bukiya, S. Durdagi, S. Noskov, A. Rosenhouse-Dantsker, Cholesterol up-regulates neuronal G protein-gated inwardly rectifying potassium (GIRK) channel activity in the hippocampus, J Biol Chem, 292 (2017) 6135–6147.

[62] C. Liu, I.S. Chen, M. Tateyama, Y. Kubo, Structural determinants of the direct inhibition of GIRK channels by Sigma-1 receptor antagonist, J Biol Chem, 300 (2024) 107219.

[63] I.S. Chen, M. Tateyama, Y. Fukata, M. Uesugi, Y. Kubo, Ivermectin activates GIRK channels in a PIP(2) −dependent, G(βγ) −independent manner and an amino acid residue at the slide helix governs the activation, J Physiol, 595 (2017) 5895–5912.

[64] A. Bertran-Mostazo, G. Putriūtė, I. Álvarez-Berbel, M.A. Busquets, C. Galdeano, A. Espargaró, R. Sabate, Proximity-Induced Pharmacology for Amyloid-Related Diseases, Cells, 13 (2024).

[65] T. Tikka, T. Usenius, M. Tenhunen, R. Keinänen, J. Koistinaho, Tetracycline derivatives and ceftriaxone, a cephalosporin antibiotic, protect neurons against apoptosis induced by ionizing radiation, J Neurochem, 78 (2001) 1409–1414.

[66] D.V. Amakhin, E.B. Soboleva, A.V. Zaitsev, Cephalosporin antibiotics are weak blockers of GABAa receptor-mediated synaptic transmission in rat brain slices, Biochem Biophys Res Commun, 499 (2018) 868–874.

[67] P. Wanleenuwat, N. Suntharampillai, P. Iwanowski, Antibiotic-induced epileptic seizures: mechanisms of action and clinical considerations, Seizure, 81 (2020) 167–174.

[68] C. Lacroix, F. Kheloufi, F. Montastruc, Y. Bennis, V. Pizzoglio, J. Micallef, Serious central nervous system side effects of cephalosporins: A national analysis of serious reports registered in the French Pharmacovigilance Database, J Neurol Sci, 398 (2019) 196–201.

[69] C. Lacroix, A.P. Bera-Jonville, F. Montastruc, L. Velly, J. Micallef, R. Guilhaumou, Serious Neurological Adverse Events of Ceftriaxone, Antibiotics (Basel), 10 (2021).

[70] J.D. Bergstrom, M.M. Kurtz, D.J. Rew, A.M. Amend, J.D. Karkas, R.G. Bostedor, V.S. Bansal, C. Dufresne, F.L. VanMiddlesworth, O.D. Hensens, et al., Zaragozic acids: a family of fungal metabolites that are picomolar competitive inhibitors of squalene synthase, Proc Natl Acad Sci U S A, 90 (1993) 80–84.

[71] R. Do, R.S. Kiss, D. Gaudet, J.C. Engert, Squalene synthase: a critical enzyme in the cholesterol biosynthesis pathway, Clin Genet, 75 (2009) 19–29.

[72] M.S. Jeeyavudeen, J.M. Pappachan, G. Arunagirinathan, Statin-related Muscle Toxicity: An Evidence-based Review, touchREV Endocrinol, 18 (2022) 89–95.

[73] L.D. Averbukh, A. Turshudzhyan, D.C. Wu, G.Y. Wu, Statin-induced Liver Injury Patterns: A Clinical Review, J Clin Transl Hepatol, 10 (2022) 543–552.

[74] S. Duddu, Y.T. Katakia, R. Chakrabarti, P. Sharma, P.C. Shukla, New epigenome players in the regulation of PCSK9-H3K4me3 and H3K9ac alterations by statin in hypercholesterolemia, J Lipid Res, 66 (2025) 100699.

[75] E.L. Spoiala, E. Cinteza, R. Vatasescu, M.V. Vlaiculescu, S.M. Moisa, Statins-Beyond Their Use in Hypercholesterolemia: Focus on the Pediatric Population, Children (Basel), 11 (2024).

[76] E. Olmastroni, G. Molari, N. De Beni, O. Colpani, F. Galimberti, M. Gazzotti, A. Zambon, A.L. Catapano, M. Casula, Statin use and risk of dementia or Alzheimer’s disease: a systematic review and meta-analysis of observational studies, Eur J Prev Cardiol, 29 (2022) 804–814.

[77] M. Laakso, L. Fernandes Silva, Statins and risk of type 2 diabetes: mechanism and clinical implications, Front Endocrinol (Lausanne), 14 (2023) 1239335.

[78] N. Ricco, S.J. Kron, Statins in Cancer Prevention and Therapy, Cancers (Basel), 15 (2023).

[79] A.H. Tremoulet, The role of statins in inflammatory vasculitides, Autoimmunity, 48 (2015) 177–180.

[80] A.N. Bukiya, P.S. Blank, A. Rosenhouse-Dantsker, Cholesterol intake and statin use regulate neuronal G protein-gated inwardly rectifying potassium channels, J Lipid Res, 60 (2019) 19–29.

[81] A. Teisseyre, A. Uryga, K. Michalak, Statins as inhibitors of voltage-gated potassium channels Kv1.3 in cancer cells, J Mol Struct, 1230 (2021) 129905.

[82] A. Teisseyre, M. Chmielarz, A. Uryga, K. Środa-Pomianek, A. Palko-Łabuz, Co-Application of Statin and Flavonoids as an Effective Strategy to Reduce the Activity of Voltage-Gated Potassium Channels Kv1.3 and Induce Apoptosis in Human Leukemic T Cell Line Jurkat, Molecules, 27 (2022).

[83] W. Ali, N. Ali, A. Ullah, S.U. Rahman, S. Ahmad, Pitavastatin and Lovastatin Exhibit Calcium Channel Blocking Activity Which Potentiate Vasorelaxant Effects of Amlodipine: A New Futuristic Dimension in Statin’s Pleiotropy, Medicina (Kaunas), 59 (2023).

[84] F. Zerin, N. Hoque, S.N. Menon, E. Ezewudo, N.P. Simon, S. Sooreni, M.S. Shahid, M. Jones, A. Pandey, Y. Gökçe, T. Rahman, R. Hasan, Nanomolar therapeutic concentrations of statins rapidly induce cerebral artery vasoconstriction by stimulating L-type calcium channels, Biochem Pharmacol, 238 (2025) 116970.

[85] M.R. Wenk, L. Lucast, G. Di Paolo, A.J. Romanelli, S.F. Suchy, R.L. Nussbaum, G.W. Cline, G.I. Shulman, W. McMurray, P. De Camilli, Phosphoinositide profiling in complex lipid mixtures using electrospray ionization mass spectrometry, Nat Biotechnol, 21 (2003) 813–817.

[86] K. D’Souza, R.M. Epand, Enrichment of phosphatidylinositols with specific acyl chains, Biochim Biophys Acta, 1838 (2014) 1501–1508.

[87] A.M. Hicks, C.J. DeLong, M.J. Thomas, M. Samuel, Z. Cui, Unique molecular signatures of glycerophospholipid species in different rat tissues analyzed by tandem mass spectrometry, Biochim Biophys Acta, 1761 (2006) 1022–1029.

[88] M. Haag, A. Schmidt, T. Sachsenheimer, B. Brügger, Quantification of Signaling Lipids by Nano-Electrospray Ionization Tandem Mass Spectrometry (Nano-ESI MS/MS), Metabolites, 2 (2012) 57–76.

[89] M. Rapedius, M. Soom, E. Shumilina, D. Schulze, R. Schönherr, C. Kirsch, F. Lang, S.J. Tucker, T. Baukrowitz, Long chain CoA esters as competitive antagonists of phosphatidylinositol 4,5-bisphosphate activation in Kir channels, J Biol Chem, 280 (2005) 30760–30767.

[90] S.B. Hansen, Lipid agonism: The PIP2 paradigm of ligand-gated ion channels, Biochim Biophys Acta, 1851 (2015) 620–628.

[91] B. Hille, E.J. Dickson, M. Kruse, O. Vivas, B.C. Suh, Phosphoinositides regulate ion channels, Biochim Biophys Acta, 1851 (2015) 844–856.

[92] B.H. Lee, J.J. De Jesús Pérez, V. Moiseenkova-Bell, T. Rohacs, Structural basis of the activation of TRPV5 channels by long-chain acyl-Coenzyme-A, Nat Commun, 14 (2023) 5883.

[93] R. Bra□nstro□m, B.E. Corkey, P.-O. Berggren, O. Larsson, Evidence for a unique long chain acyl-CoA ester binding site on the ATP-regulated potassium channel in mouse pancreatic beta cells, Journal of Biological Chemistry, 272 (1997) 17390–17394.

[94] C.L. Huang, S. Feng, D.W. Hilgemann, Direct activation of inward rectifier potassium channels by PIP2 and its stabilization by Gbetagamma, Nature, 391 (1998) 803–806.

[95] J. Han, D. Kang, D. Kim, Properties and modulation of the G protein-coupled K+ channel in rat cerebellar granule neurons: ATP versus phosphatidylinositol 4,5-bisphosphate, J Physiol, 550 (2003) 693–706.

[96] C.A. Calderón-Ospina, M.O. Nava-Mesa, B Vitamins in the nervous system: Current knowledge of the biochemical modes of action and synergies of thiamine, pyridoxine, and cobalamin, CNS Neurosci Ther, 26 (2020) 5–13.

[97] C.K. Singleton, P.R. Martin, Molecular mechanisms of thiamine utilization, Curr Mol Med, 1 (2001) 197–207.

[98] A. Bâ, Metabolic and structural role of thiamine in nervous tissues, Cell Mol Neurobiol, 28 (2008) 923–931.

[99] A. Di Gennaro, J.Z. Haeggström, The leukotrienes: immune-modulating lipid mediators of disease, Adv Immunol, 116 (2012) 51–92.

[100] V. Sharma, P. Sharma, T.G. Singh, Leukotriene signaling in neurodegeneration: implications for treatment strategies, Inflammopharmacology, 32 (2024) 3571–3584.

[101] D. Kim, D.L. Lewis, L. Graziadei, E.J. Neer, D. Bar-Sagi, D.E. Clapham, G-protein beta gamma-subunits activate the cardiac muscarinic K+-channel via phospholipase A2, Nature, 337 (1989) 557–560.

[102] Y. Kurachi, H. Ito, T. Sugimoto, T. Shimizu, I. Miki, M. Ui, Arachidonic acid metabolites as intracellular modulators of the G protein-gated cardiac K+ channel, Nature, 337 (1989) 555–557.

[103] R.W. Scherer, C.F. Lo, G.E. Breitwieser, Leukotriene C4 modulation of muscarinic K+ current activation in bullfrog atrial myocytes, J Gen Physiol, 102 (1993) 125–141.

[104] A.N. Bukiya, J. McMillan, J. Liu, B. Shivakumar, A.L. Parrill, A.M. Dopico, Activation of calcium- and voltage-gated potassium channels of large conductance by leukotriene B4, J Biol Chem, 289 (2014) 35314–35325.

[105] S. Bang, S. Yoo, T.J. Yang, H. Cho, S.W. Hwang, 17(R)-resolvin D1 specifically inhibits transient receptor potential ion channel vanilloid 3 leading to peripheral antinociception, Br J Pharmacol, 165 (2012) 683–692.

[106] F. Quan-Xin, F. Fan, F. Xiang-Ying, L. Shu-Jun, W. Shi-Qi, L. Zhao-Xu, Z. Xu-Jie, Z. Qing-Chuan, W. Wei, Resolvin D1 reverses chronic pancreatitis-induced mechanical allodynia, phosphorylation of NMDA receptors, and cytokines expression in the thoracic spinal dorsal horn, BMC Gastroenterol, 12 (2012) 148.

[107] Z.Z. Xu, L. Zhang, T. Liu, J.Y. Park, T. Berta, R. Yang, C.N. Serhan, R.R. Ji, Resolvins RvE1 and RvD1 attenuate inflammatory pain via central and peripheral actions, Nat Med, 16 (2010) 592–597, 591p following 597.

[108] C.K. Park, Z.Z. Xu, T. Liu, N. Lü, C.N. Serhan, R.R. Ji, Resolvin D2 is a potent endogenous inhibitor for transient receptor potential subtype V1/A1, inflammatory pain, and spinal cord synaptic plasticity in mice: distinct roles of resolvin D1, D2, and E1, J Neurosci, 31 (2011) 18433–18438.

[109] S.Y. Jeong, H.L. Lee, S. Wee, H. Lee, G. Hwang, S. Hwang, S. Yoon, Y.I. Yang, I. Han, K.N. Kim, Co-Administration of Resolvin D1 and Peripheral Nerve-Derived Stem Cell Spheroids as a Therapeutic Strategy in a Rat Model of Spinal Cord Injury, Int J Mol Sci, 24 (2023).

[110] Y. Wang, Y. Yang, S. Zhang, C. Li, L. Zhang, Modulation of neuroinflammation by cysteinyl leukotriene 1 and 2 receptors: implications for cerebral ischemia and neurodegenerative diseases, Neurobiol Aging, 87 (2020) 1–10.

[111] J. Michael, M.S. Unger, R. Poupardin, P. Schernthaner, H. Mrowetz, J. Attems, L. Aigner, Microglia depletion diminishes key elements of the leukotriene pathway in the brain of Alzheimer’s Disease mice, Acta Neuropathol Commun, 8 (2020) 129.

[112] K. Strempfl, M.S. Unger, S. Flunkert, A. Trost, H.A. Reitsamer, B. Hutter-Paier, L. Aigner, Leukotriene Signaling as a Target in α-Synucleinopathies, Biomolecules, 12 (2022).

[113] J. Michael, J. Zirknitzer, M.S. Unger, R. Poupardin, T. Rieß, N. Paiement, H. Zerbe, B. Hutter-Paier, H. Reitsamer, L. Aigner, The Leukotriene Receptor Antagonist Montelukast Attenuates Neuroinflammation and Affects Cognition in Transgenic 5xFAD Mice, Int J Mol Sci, 22 (2021).

[114] S. Chamani, V. Bianconi, A. Tasbandi, M. Pirro, G.E. Barreto, T. Jamialahmadi, A. Sahebkar, Resolution of Inflammation in Neurodegenerative Diseases: The Role of Resolvins, Mediators Inflamm, 2020 (2020) 3267172.

[115] M.T. Mizwicki, G. Liu, M. Fiala, L. Magpantay, J. Sayre, A. Siani, M. Mahanian, R. Weitzman, E.Y. Hayden, M.J. Rosenthal, I. Nemere, J. Ringman, D.B. Teplow, 1α,25-dihydroxyvitamin D3 and resolvin D1 retune the balance between amyloid-β phagocytosis and inflammation in Alzheimer’s disease patients, J Alzheimers Dis, 34 (2013) 155–170.

[116] P. Krashia, A. Cordella, A. Nobili, L. La Barbera, M. Federici, A. Leuti, F. Campanelli, G. Natale, G. Marino, V. Calabrese, F. Vedele, V. Ghiglieri, B. Picconi, G. Di Lazzaro, T. Schirinzi, G. Sancesario, N. Casadei, O. Riess, S. Bernardini, A. Pisani, P. Calabresi, M.T. Viscomi, C.N. Serhan, V. Chiurchiù, M. D’Amelio, N.B. Mercuri, Blunting neuroinflammation with resolvin D1 prevents early pathology in a rat model of Parkinson’s disease, Nat Commun, 10 (2019) 3945.

[117] C.F. Marques, M.M. Marques, G.C. Justino, The mechanisms underlying montelukast’s neuropsychiatric effects - new insights from a combined metabolic and multiomics approach, Life Sci, 310 (2022) 121056.

[118] S. Attaluri, R. Upadhya, M. Kodali, L.N. Madhu, D. Upadhya, B. Shuai, A.K. Shetty, Brain-Specific Increase in Leukotriene Signaling Accompanies Chronic Neuroinflammation and Cognitive Impairment in a Model of Gulf War Illness, Front Immunol, 13 (2022) 853000.

[119] C. Li, X. Wu, S. Liu, D. Shen, J. Zhu, K. Liu, Role of Resolvins in the Inflammatory Resolution of Neurological Diseases, Front Pharmacol, 11 (2020) 612.

[120] B. Kok Kendirlioglu, P. Unalan Ozpercin, O. Yuksel Oksuz, S. Sozen, R. Cihnioglu, T. Kalelioglu, M.C. Ilnem, N. Karamustafalioglu, Resolvin D1 as a novel anti-inflammatory marker in manic, depressive and euthymic states of bipolar disorder, Nord J Psychiatry, 74 (2020) 83–88.

[121] N.L. Del Rey, N. Hernández-Pinedo, M. Carrillo, M. Del Cerro, N. Esteban-García, I. Trigo-Damas, M.H.G. Monje, J.L. Lanciego, C. Cavada, J.A. Obeso, J. Blesa, Calbindin and Girk2/Aldh1a1 define resilient vs vulnerable dopaminergic neurons in a primate Parkinson’s disease model, NPJ Parkinsons Dis, 10 (2024) 165.

[122] R. Alfaro-Ruiz, A. Martín-Belmonte, C. Aguado, F. Hernández, A.E. Moreno-Martínez, J. Ávila, R. Luján, The Expression and Localisation of G-Protein-Coupled Inwardly Rectifying Potassium (GIRK) Channels Is Differentially Altered in the Hippocampus of Two Mouse Models of Alzheimer’s Disease, Int J Mol Sci, 22 (2021).

[123] G. Choi, S.W. Hwang, Modulation of the Activities of Neuronal Ion Channels by Fatty Acid-Derived Pro-Resolvents, Front Physiol, 7 (2016) 523.

[124] M. Fiala, N. Terrando, J. Dalli, Specialized Pro-Resolving Mediators from Omega-3 Fatty Acids Improve Amyloid-β Phagocytosis and Regulate Inflammation in Patients with Minor Cognitive Impairment, J Alzheimers Dis, 48 (2015) 293–301.

[125] D.F. Baggio, F.M.R. da Luz, R.V. Lopes, L.E.N. Ferreira, E.I. Araya, J.G. Chichorro, Sex Dimorphism in Resolvin D5-induced Analgesia in Rat Models of Trigeminal Pain, J Pain, 24 (2023) 717–729.

[126] J. Park, J. Roh, J. Pan, Y.H. Kim, C.K. Park, Y.Y. Jo, Role of Resolvins in Inflammatory and Neuropathic Pain, Pharmaceuticals (Basel), 16 (2023).

[127] X. Luo, Y. Gu, X. Tao, C.N. Serhan, R.R. Ji, Resolvin D5 Inhibits Neuropathic and Inflammatory Pain in Male But Not Female Mice: Distinct Actions of D-Series Resolvins in Chemotherapy-Induced Peripheral Neuropathy, Front Pharmacol, 10 (2019) 745.

[128] J.Z. Haeggström, C.D. Funk, Lipoxygenase and leukotriene pathways: biochemistry, biology, and roles in disease, Chem Rev, 111 (2011) 5866–5898.

[129] X. Zhang, L. Chen, J.P. Hardwick, Promoter activity and regulation of the CYP4F2 leukotriene B(4) omega-hydroxylase gene by peroxisomal proliferators and retinoic acid in HepG2 cells, Arch Biochem Biophys, 378 (2000) 364–376.

[130] E. Smeets, S. Huang, X.Y. Lee, E. Van Nieuwenhove, C. Helsen, F. Handle, L. Moris, S. El Kharraz, R. Eerlings, W. Devlies, M. Willemsen, L. Bücken, T. Prezzemolo, S. Humblet-Baron, A. Voet, A. Rochtus, A. Van Schepdael, F. de Zegher, F. Claessens, A disease-associated missense mutation in CYP4F3 affects the metabolism of leukotriene B4 via disruption of electron transfer, J Cachexia Sarcopenia Muscle, 13 (2022) 2242–2253.

[131] B. Samuelsson, S.E. Dahlén, J.A. Lindgren, C.A. Rouzer, C.N. Serhan, Leukotrienes and lipoxins: structures, biosynthesis, and biological effects, Science, 237 (1987) 1171–1176.

