## Supplemental Figures S1-S26 for "Structure-based identification of GIRK2-PIP2 modulators among known drugs and metabolites using docking, MM-GBSA, ADMET, and molecular dynamics"

### Slide 1
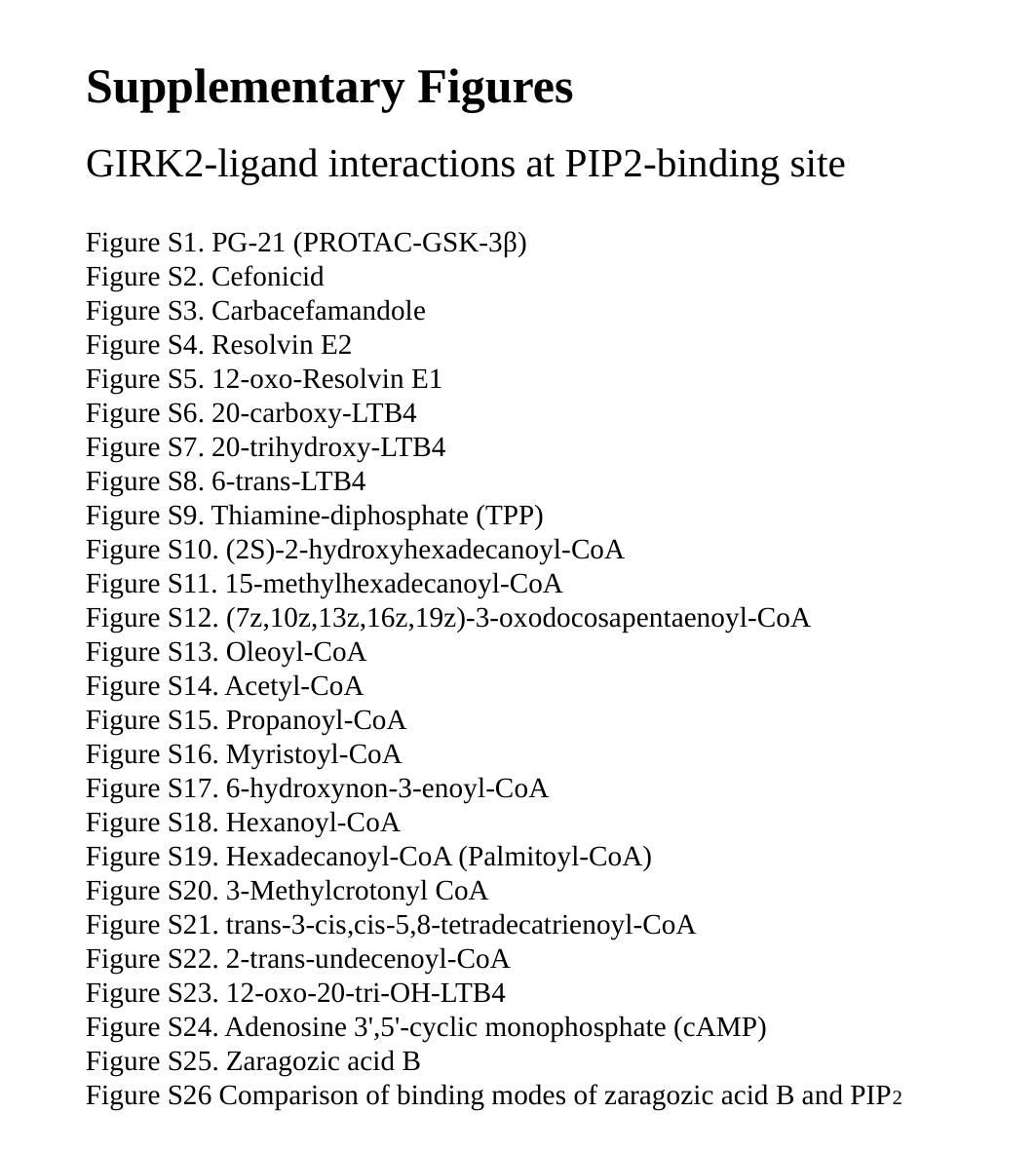

#### Supplementary Figures
GIRK2-ligand interactions at PIP2-binding site
Figure S1. PG-21 (PROTAC-GSK-3β)
Figure S2. Cefonicid
Figure S3. Carbacefamandole
Figure S4. Resolvin E2
Figure S5. 12-oxo-Resolvin E1
Figure S6. 20-carboxy-LTB4
Figure S7. 20-trihydroxy-LTB4
Figure S8. 6-trans-LTB4
Figure S9. Thiamine-diphosphate (TPP)
Figure S10. (2S)-2-hydroxyhexadecanoyl-CoA
Figure S11. 15-methylhexadecanoyl-CoA
Figure S12. (7z,10z,13z,16z,19z)-3-oxodocosapentaenoyl-CoA
Figure S13. Oleoyl-CoA
Figure S14. Acetyl-CoA
Figure S15. Propanoyl-CoA
Figure S16. Myristoyl-CoA
Figure S17. 6-hydroxynon-3-enoyl-CoA
Figure S18. Hexanoyl-CoA
Figure S19. Hexadecanoyl-CoA (Palmitoyl-CoA)
Figure S20. 3-Methylcrotonyl CoA
Figure S21. trans-3-cis,cis-5,8-tetradecatrienoyl-CoA
Figure S22. 2-trans-undecenoyl-CoA
Figure S23. 12-oxo-20-tri-OH-LTB4
Figure S24. Adenosine 3',5'-cyclic monophosphate (cAMP)
Figure S25. Zaragozic acid B
Figure S26 Comparison of binding modes of zaragozic acid B and PIP2

### Slide 2
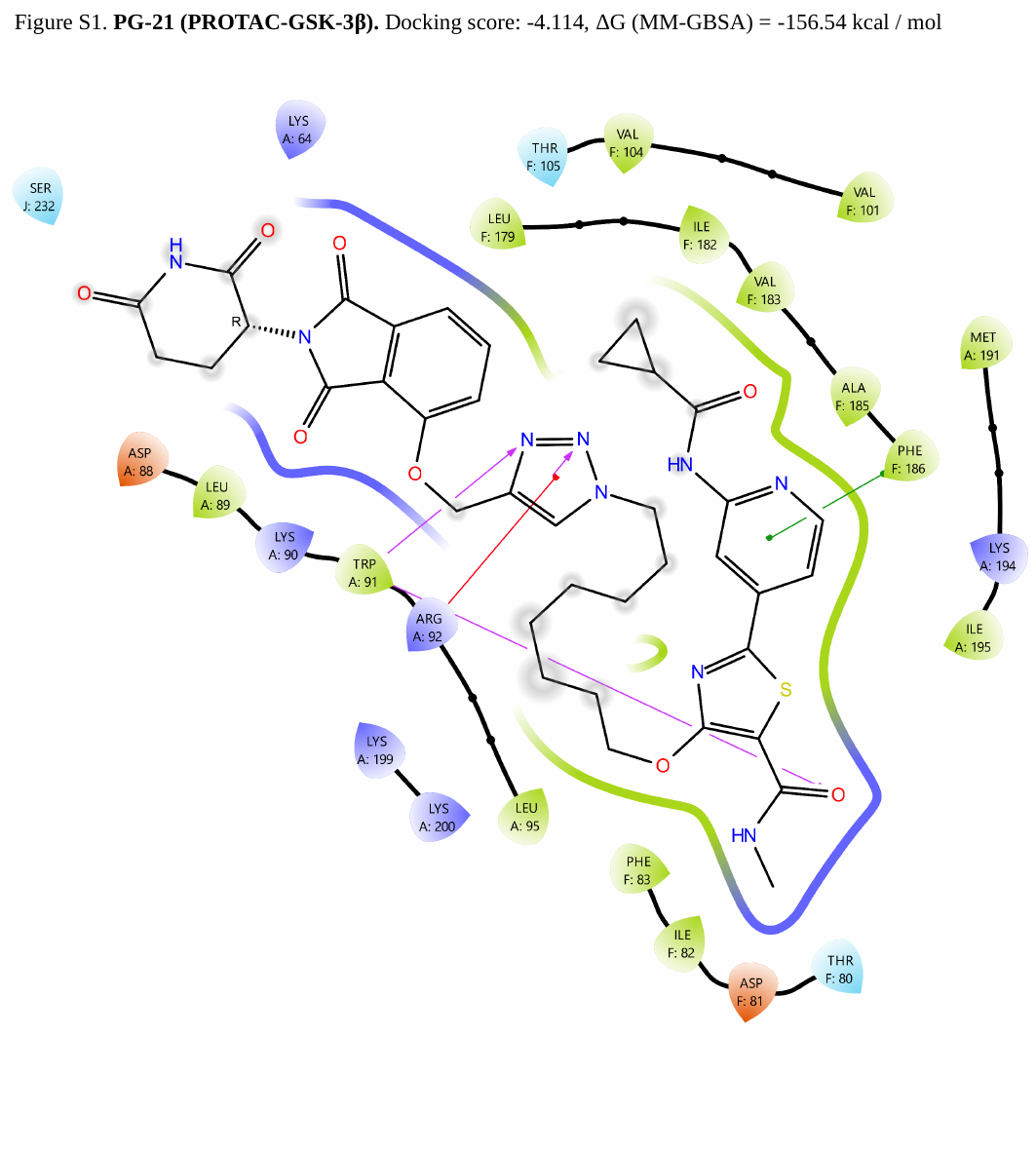

Figure S1. PG-21 (PROTAC-GSK-3β). Docking score: -4.114, ΔG (MM-GBSA) = -156.54 kcal / mol

### Slide 3
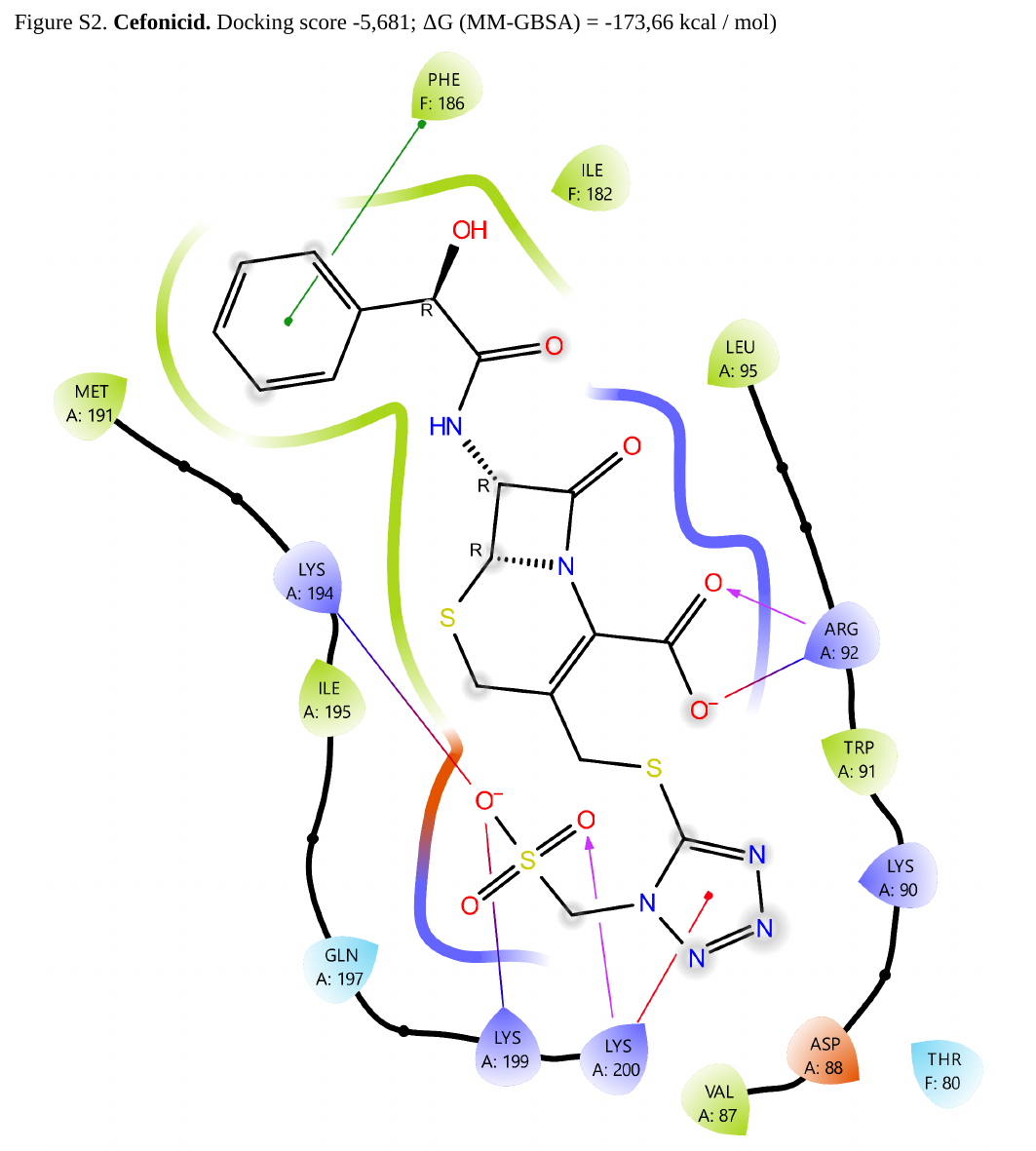

Figure S2. Cefonicid. Docking score -5,681; ΔG (MM-GBSA) = -173,66 kcal / mol)

### Slide 4
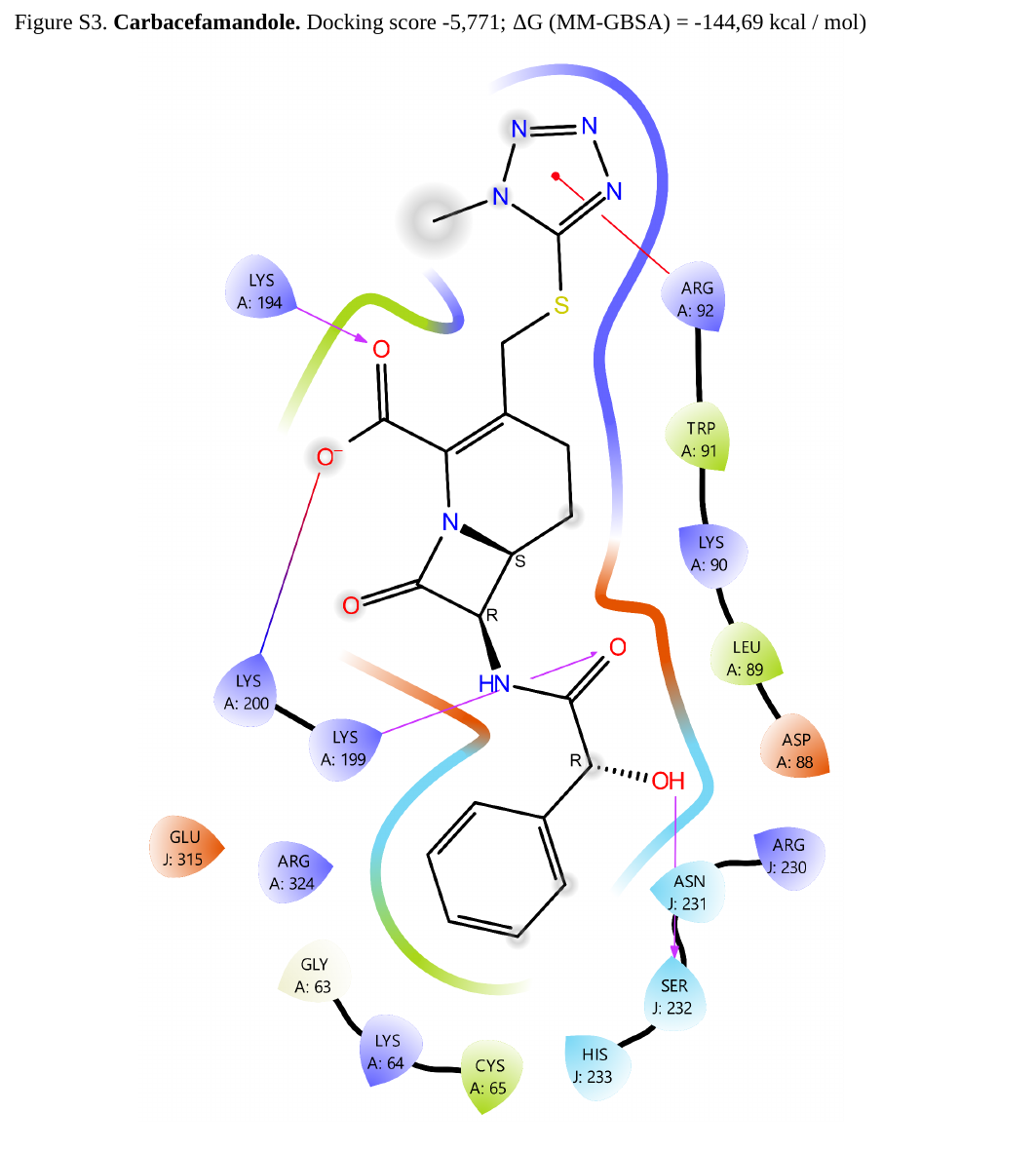

Figure S3. Carbacefamandole. Docking score -5,771; ΔG (MM-GBSA) = -144,69 kcal / mol)

### Slide 5
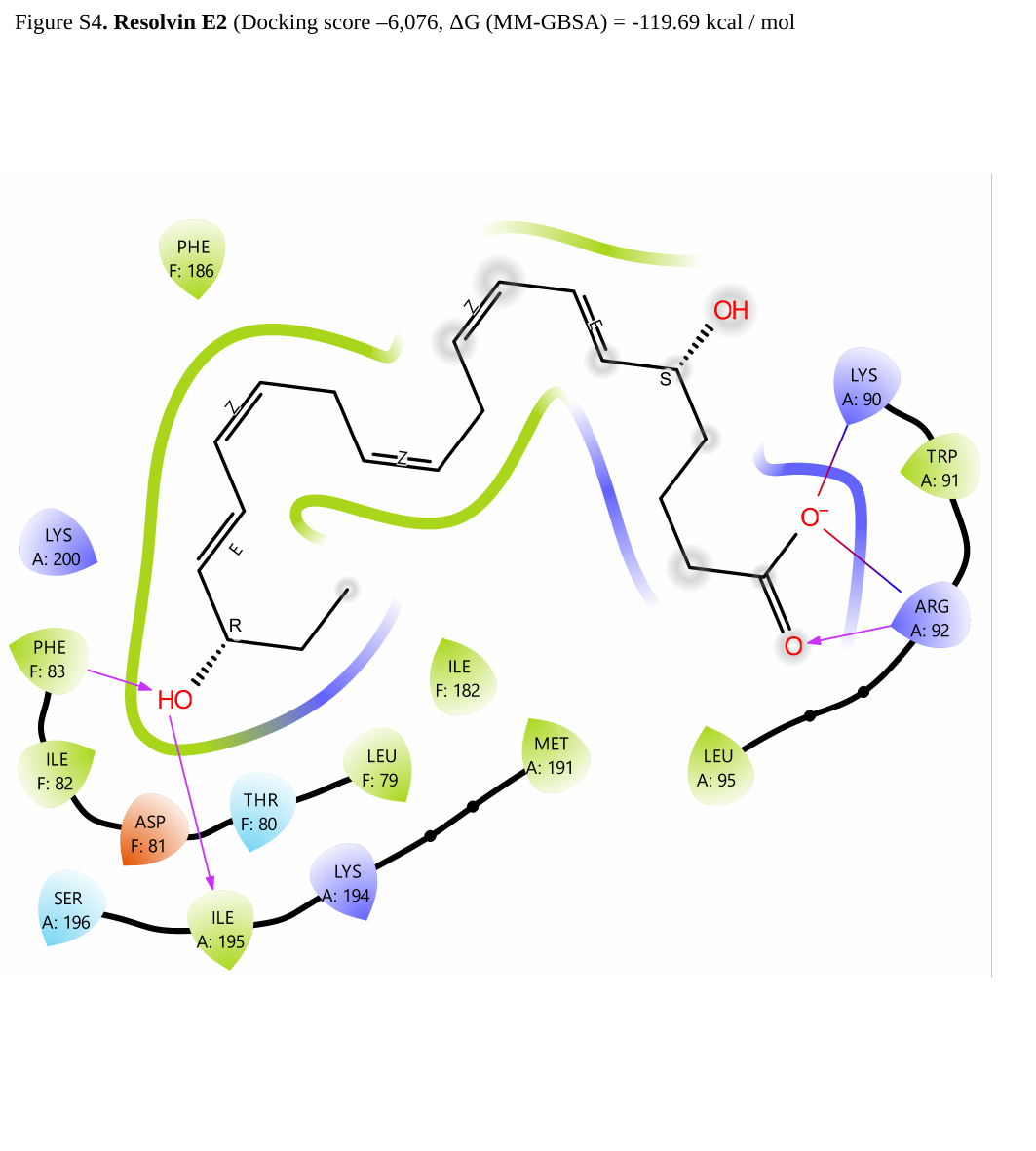

Figure S4. Resolvin E2 (Docking score –6,076, ΔG (MM-GBSA) = -119.69 kcal / mol

### Slide 6
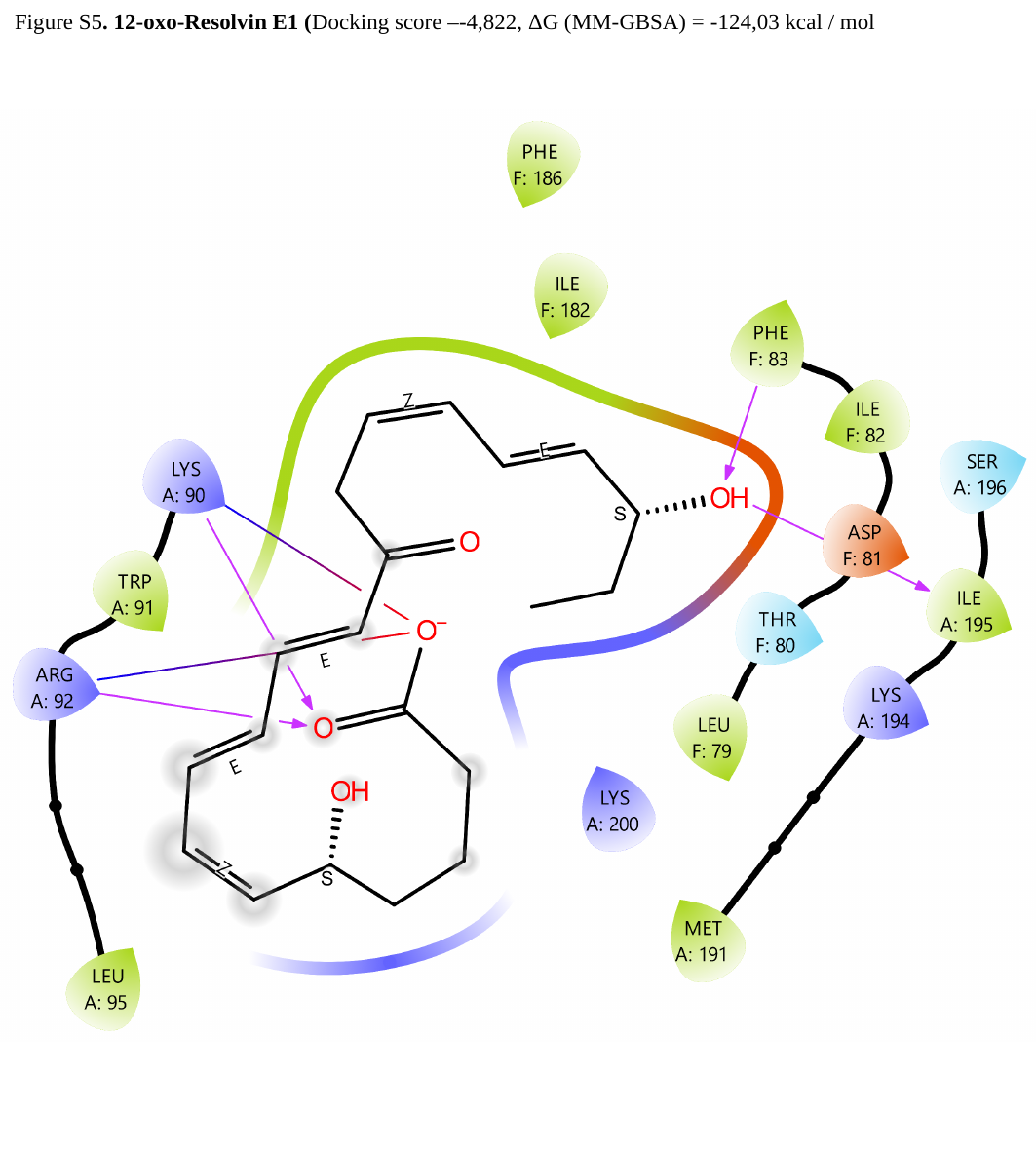

Figure S5. 12-oxo-Resolvin E1 (Docking score –-4,822, ΔG (MM-GBSA) = -124,03 kcal / mol

### Slide 7
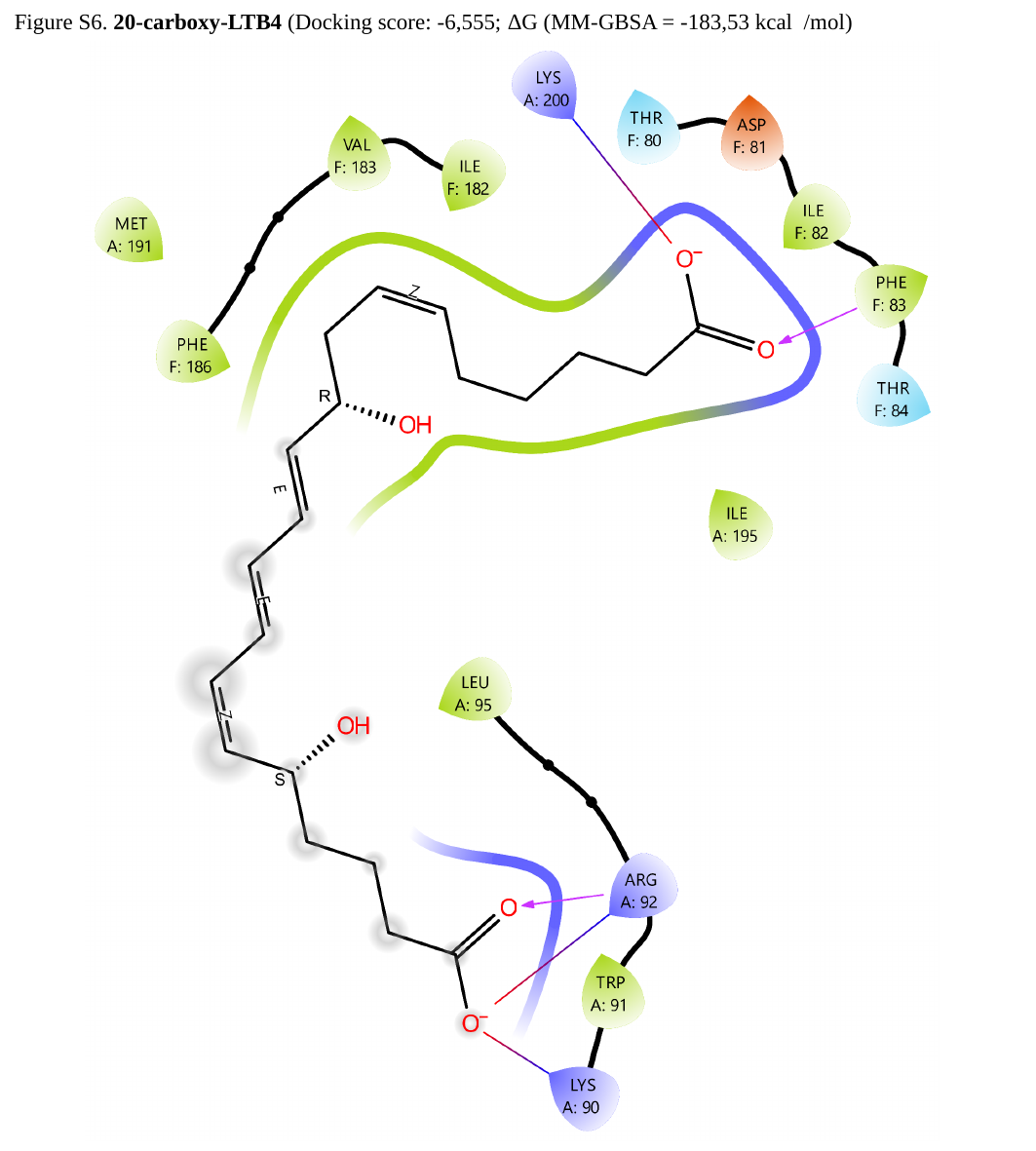

Figure S6. 20-carboxy-LTB4 (Docking score: -6,555; ΔG (MM-GBSA = -183,53 kcal /mol)

### Slide 8
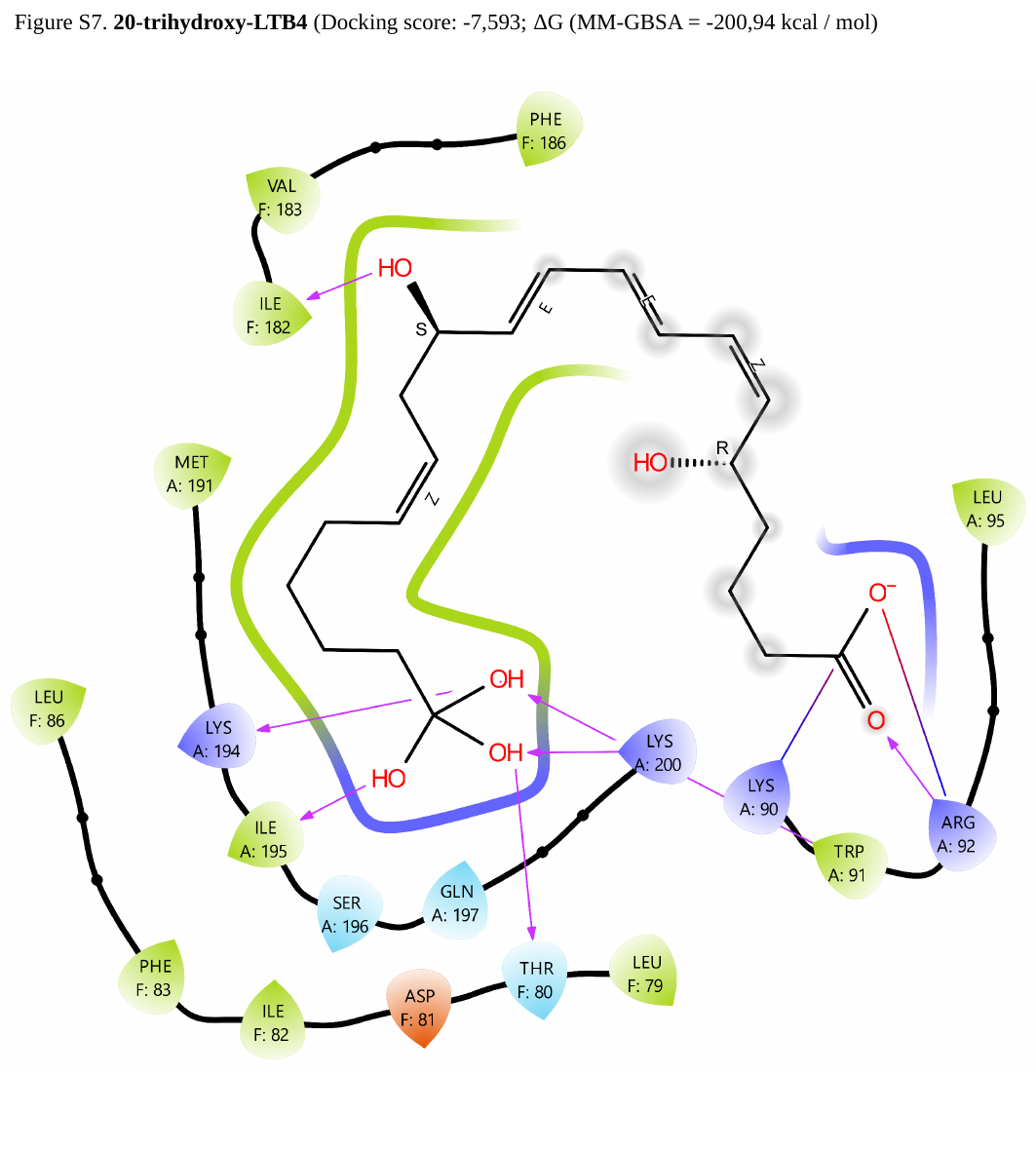

Figure S7. 20-trihydroxy-LTB4 (Docking score: -7,593; ΔG (MM-GBSA = -200,94 kcal / mol)

### Slide 9
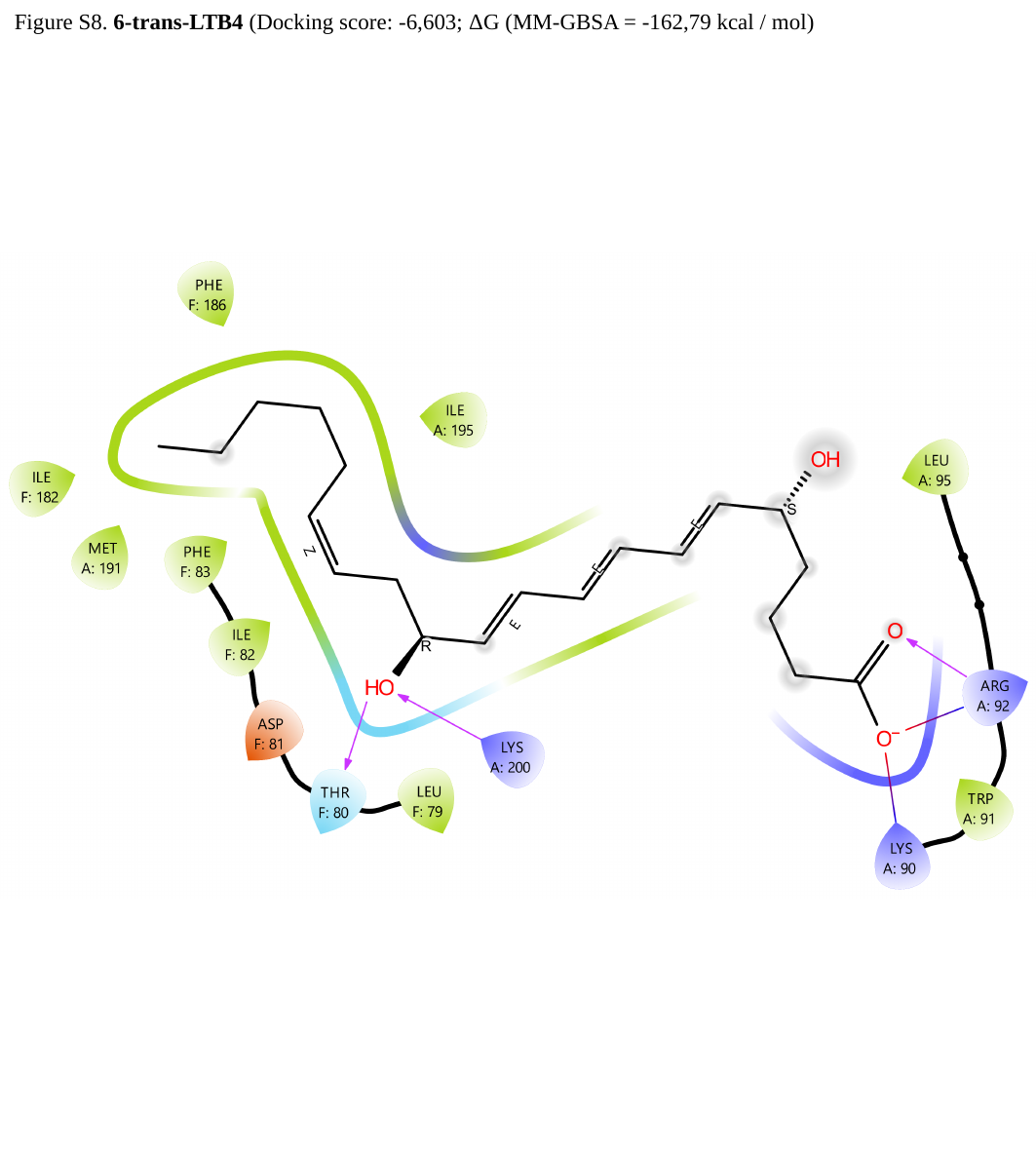

Figure S8. 6-trans-LTB4 (Docking score: -6,603; ΔG (MM-GBSA = -162,79 kcal / mol)

### Slide 10
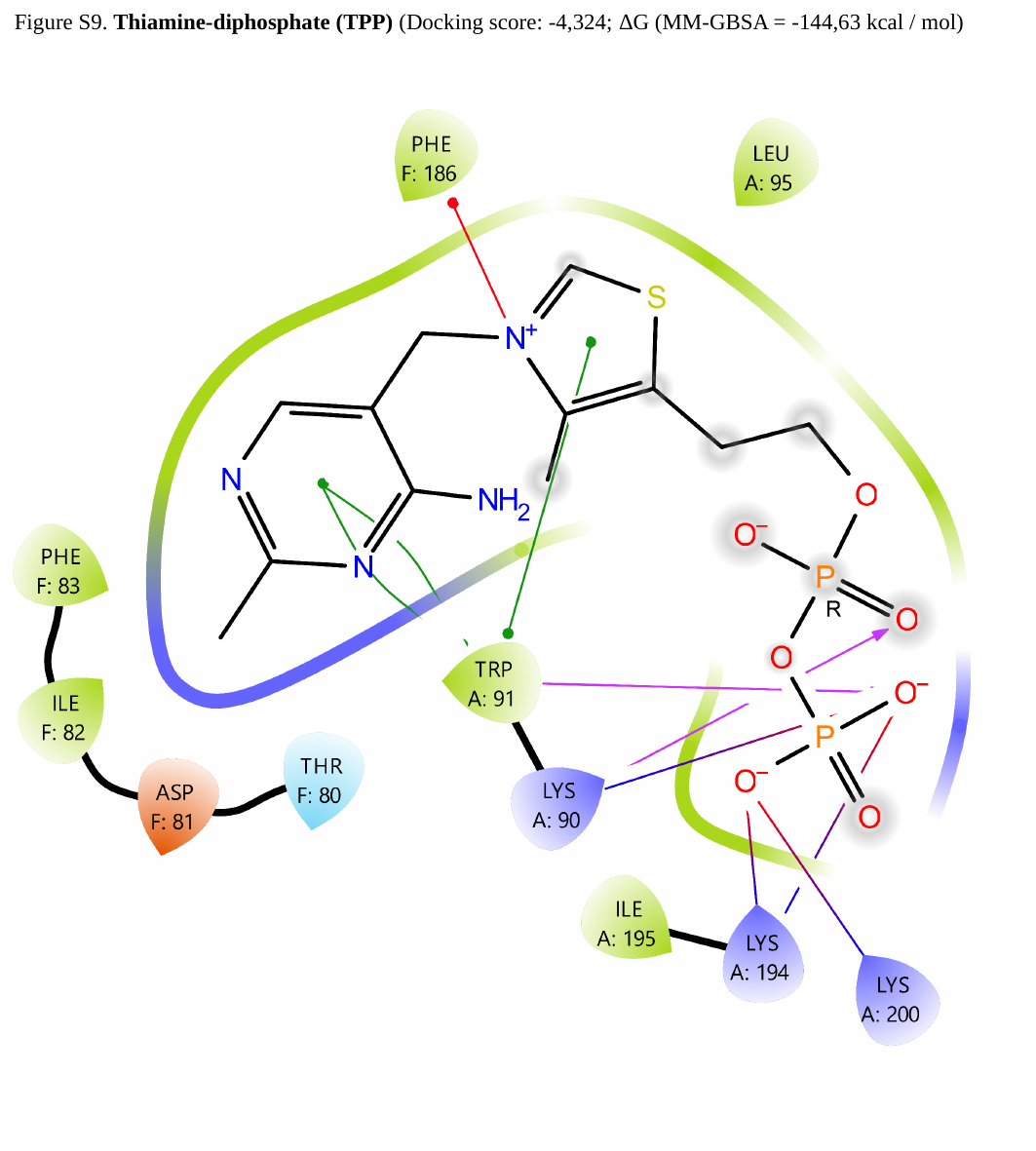

Figure S9. Thiamine-diphosphate (TPP) (Docking score: -4,324; ΔG (MM-GBSA = -144,63 kcal / mol)

### Slide 11
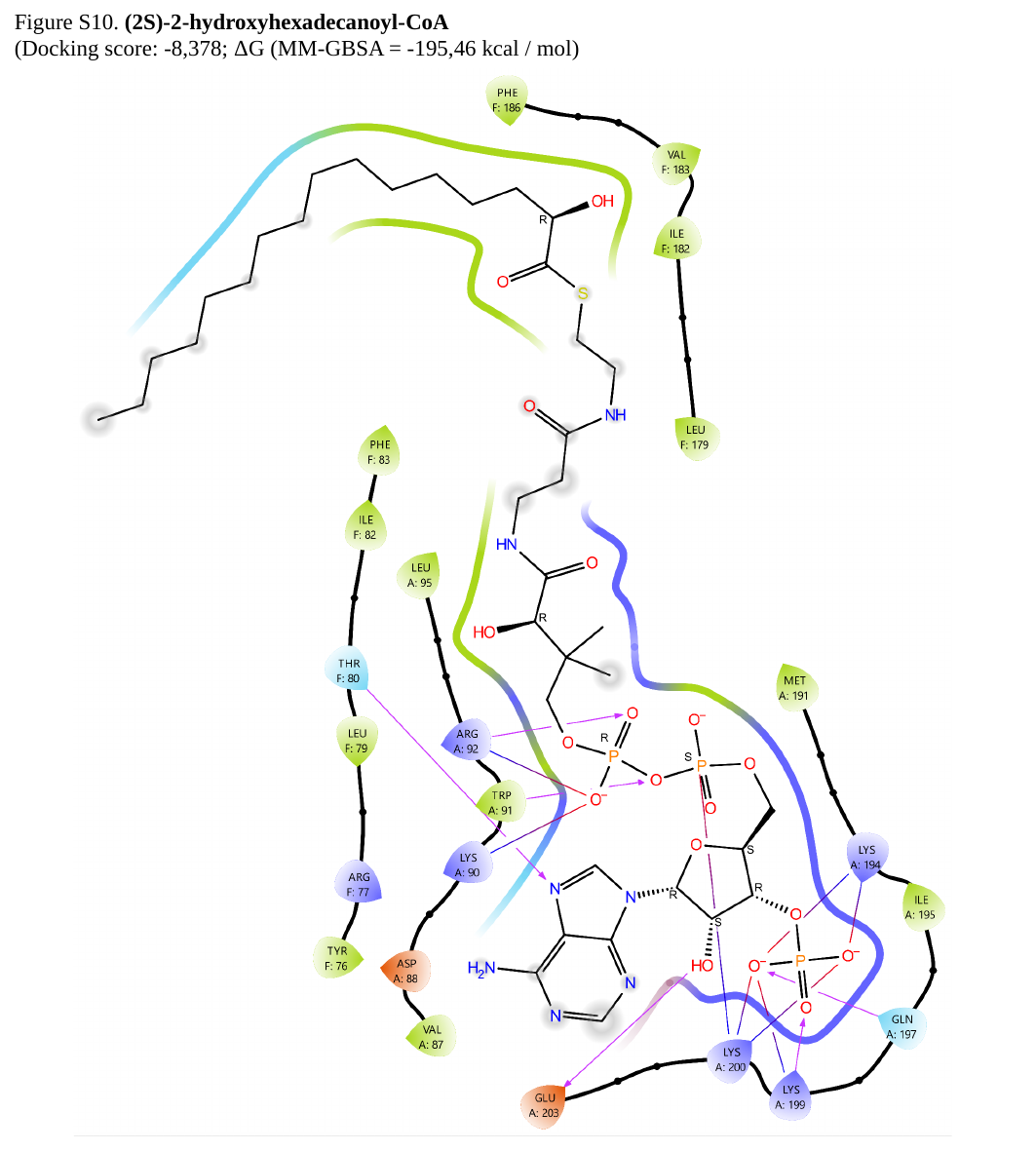

Figure S10. (2S)-2-hydroxyhexadecanoyl-CoA
(Docking score: -8,378; ΔG (MM-GBSA = -195,46 kcal / mol)

### Slide 12
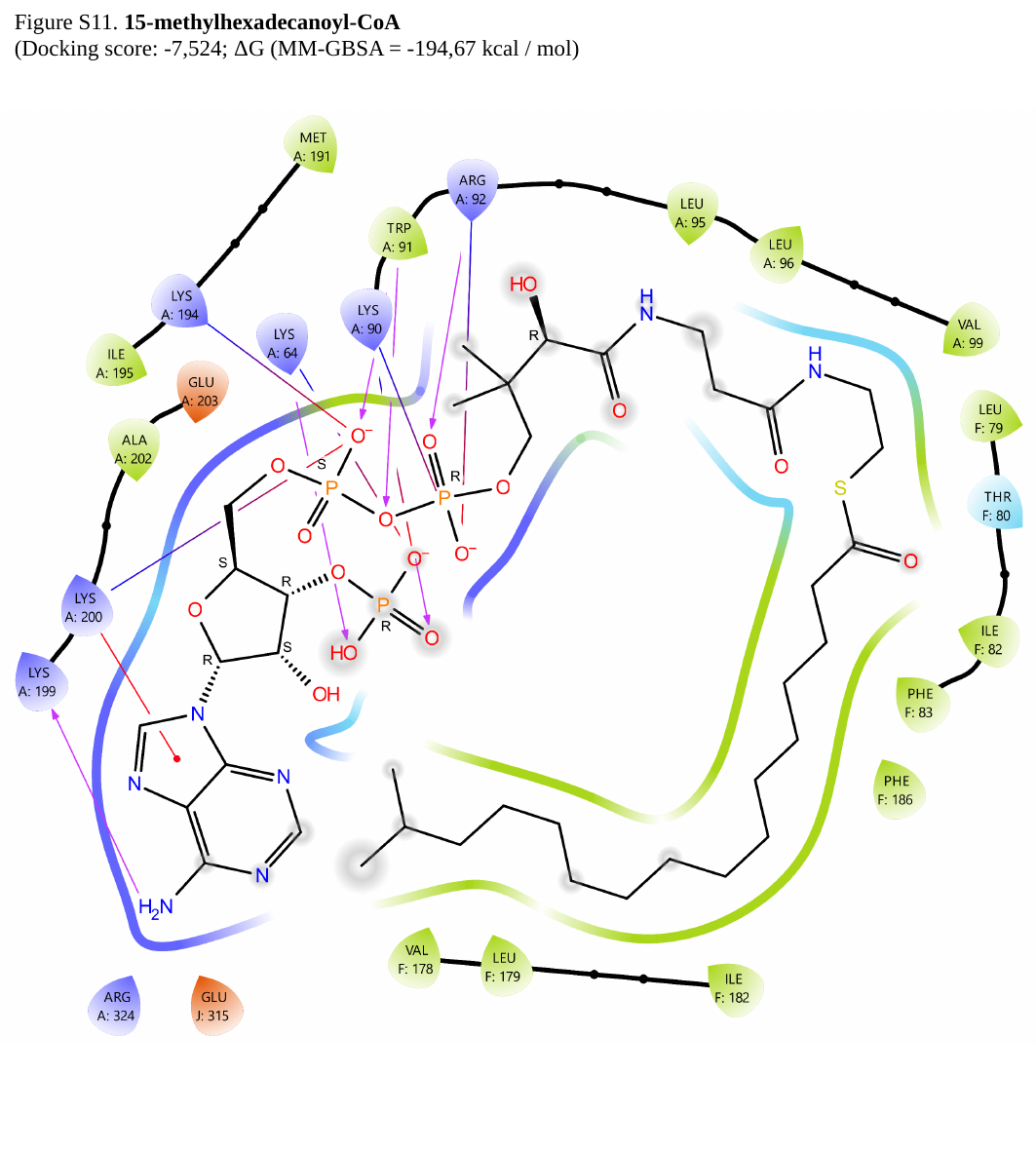

Figure S11. 15-methylhexadecanoyl-CoA
(Docking score: -7,524; ΔG (MM-GBSA = -194,67 kcal / mol)

### Slide 13
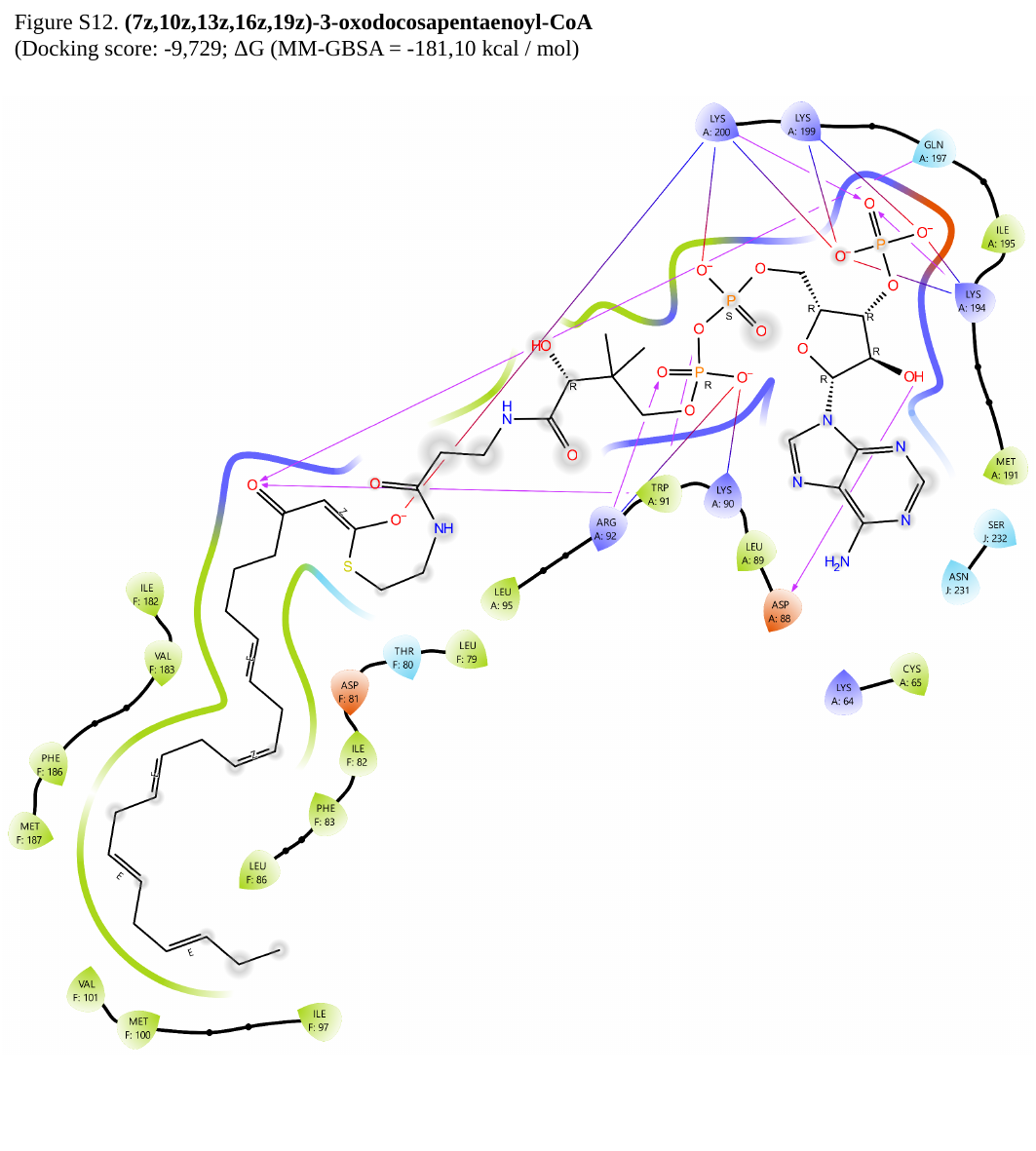

Figure S12. (7z,10z,13z,16z,19z)-3-oxodocosapentaenoyl-CoA
(Docking score: -9,729; ΔG (MM-GBSA = -181,10 kcal / mol)

### Slide 14
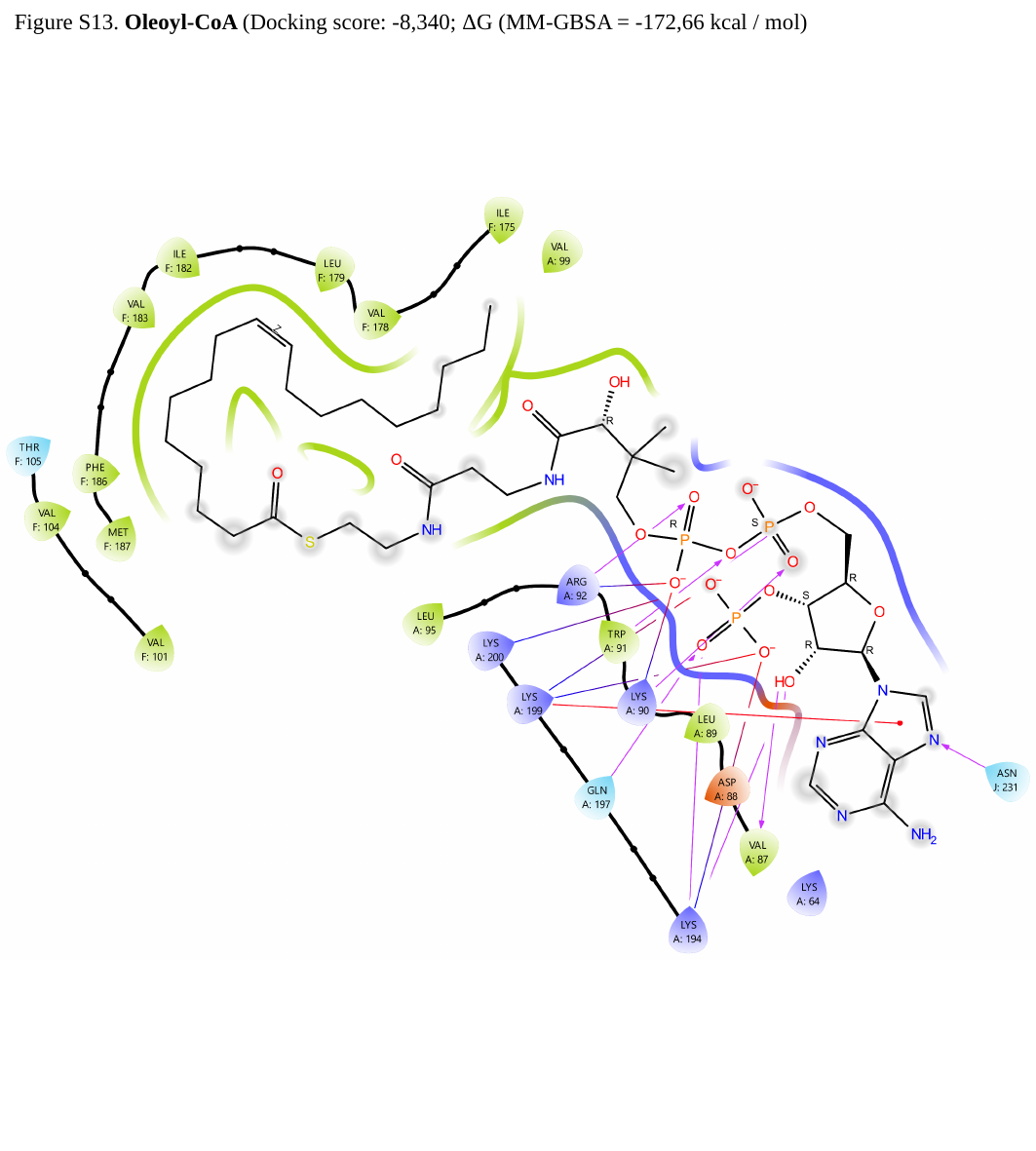

Figure S13. Oleoyl-CoA (Docking score: -8,340; ΔG (MM-GBSA = -172,66 kcal / mol)

### Slide 15
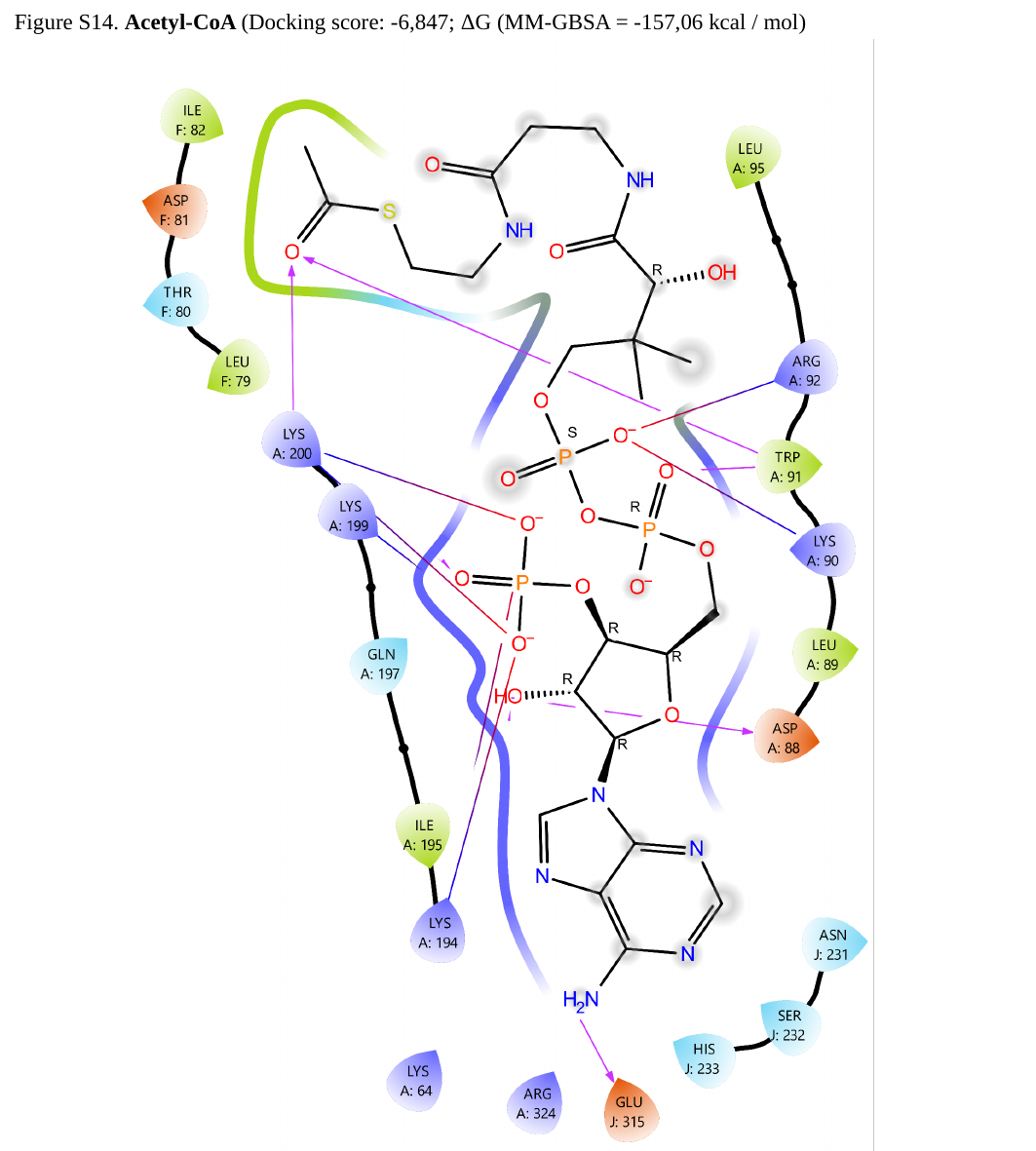

Figure S14. Acetyl-CoA (Docking score: -6,847; ΔG (MM-GBSA = -157,06 kcal / mol)

### Slide 16
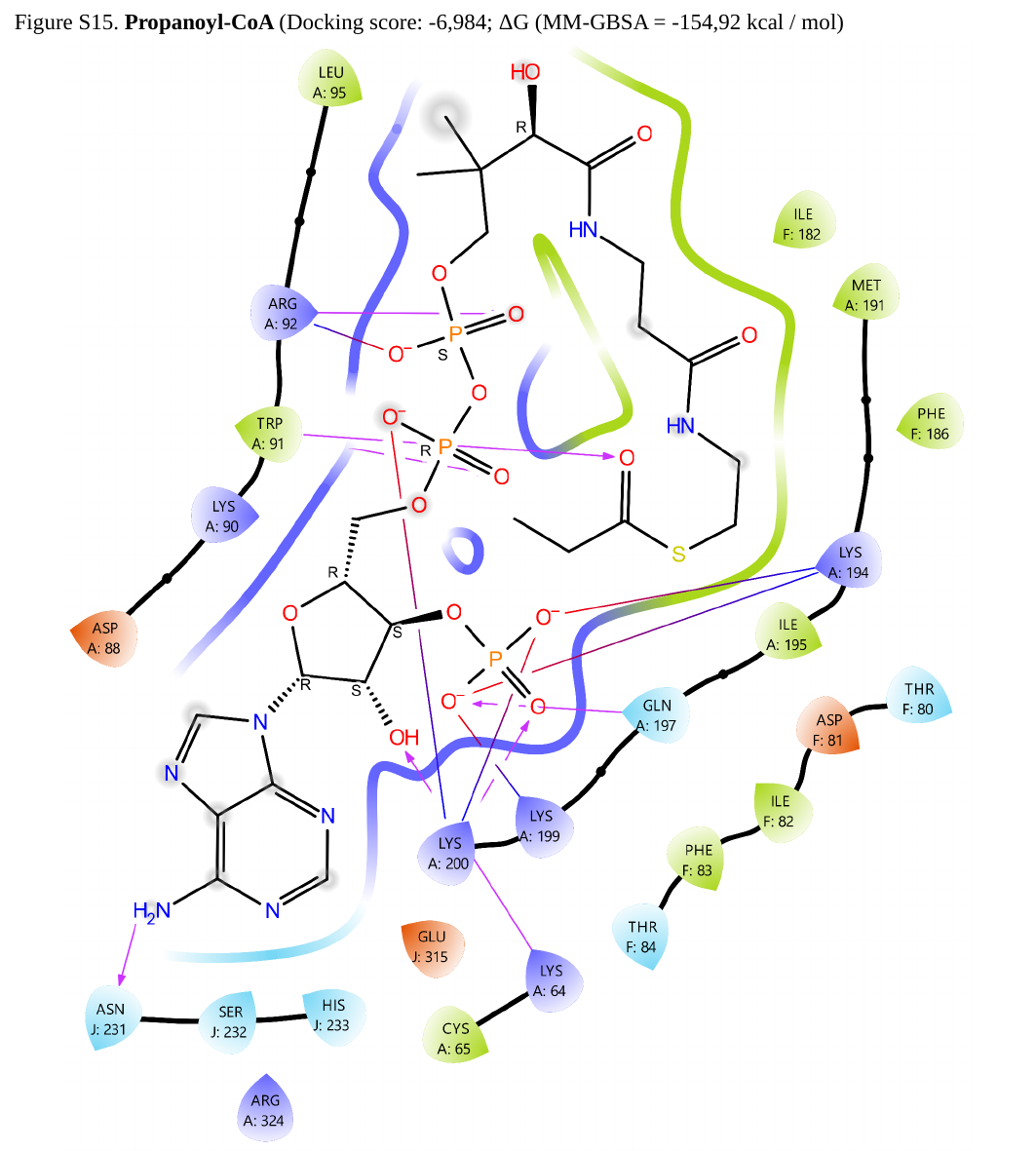

Figure S15. Propanoyl-CoA (Docking score: -6,984; ΔG (MM-GBSA = -154,92 kcal / mol)

### Slide 17
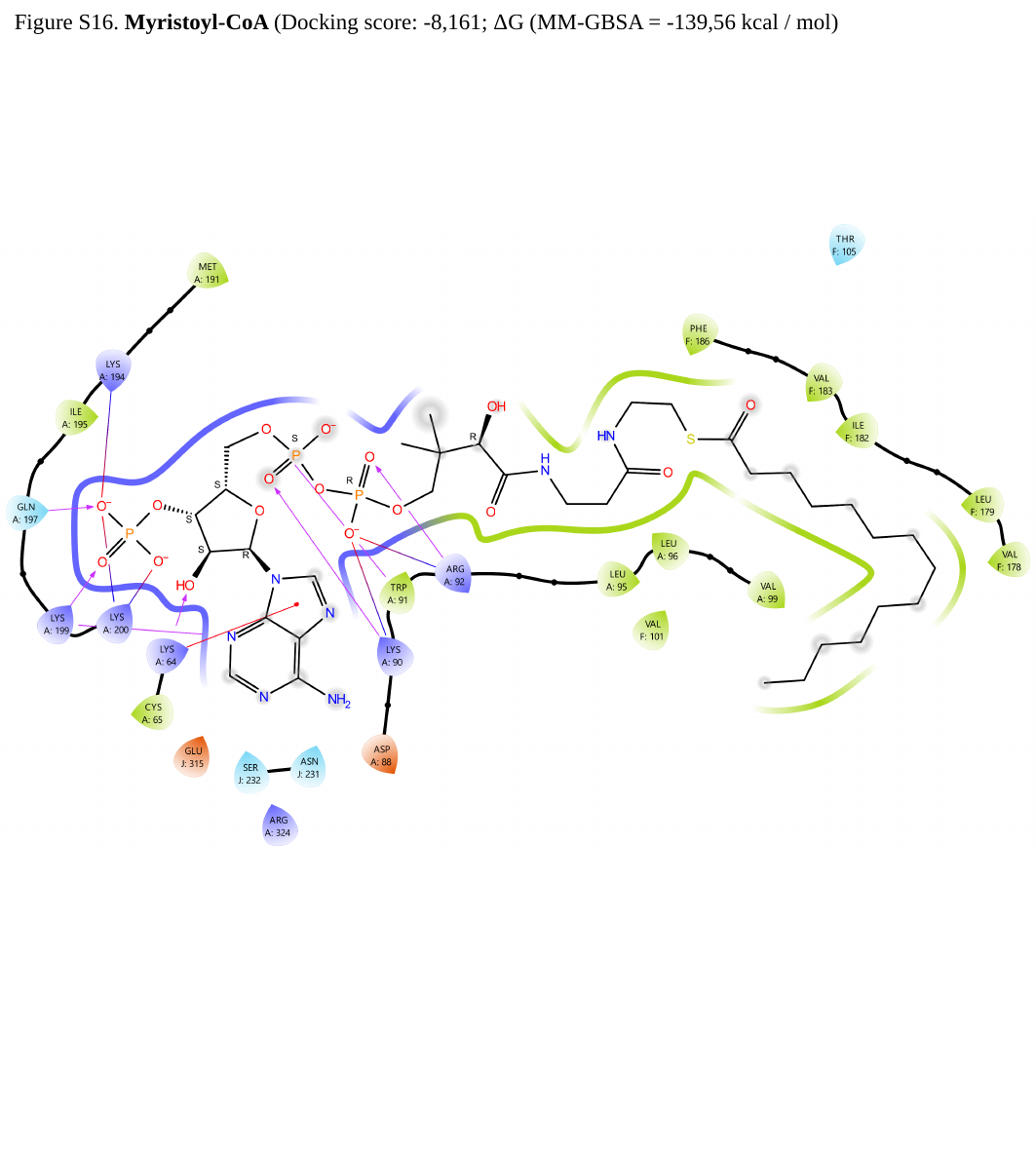

Figure S16. Myristoyl-CoA (Docking score: -8,161; ΔG (MM-GBSA = -139,56 kcal / mol)

### Slide 18
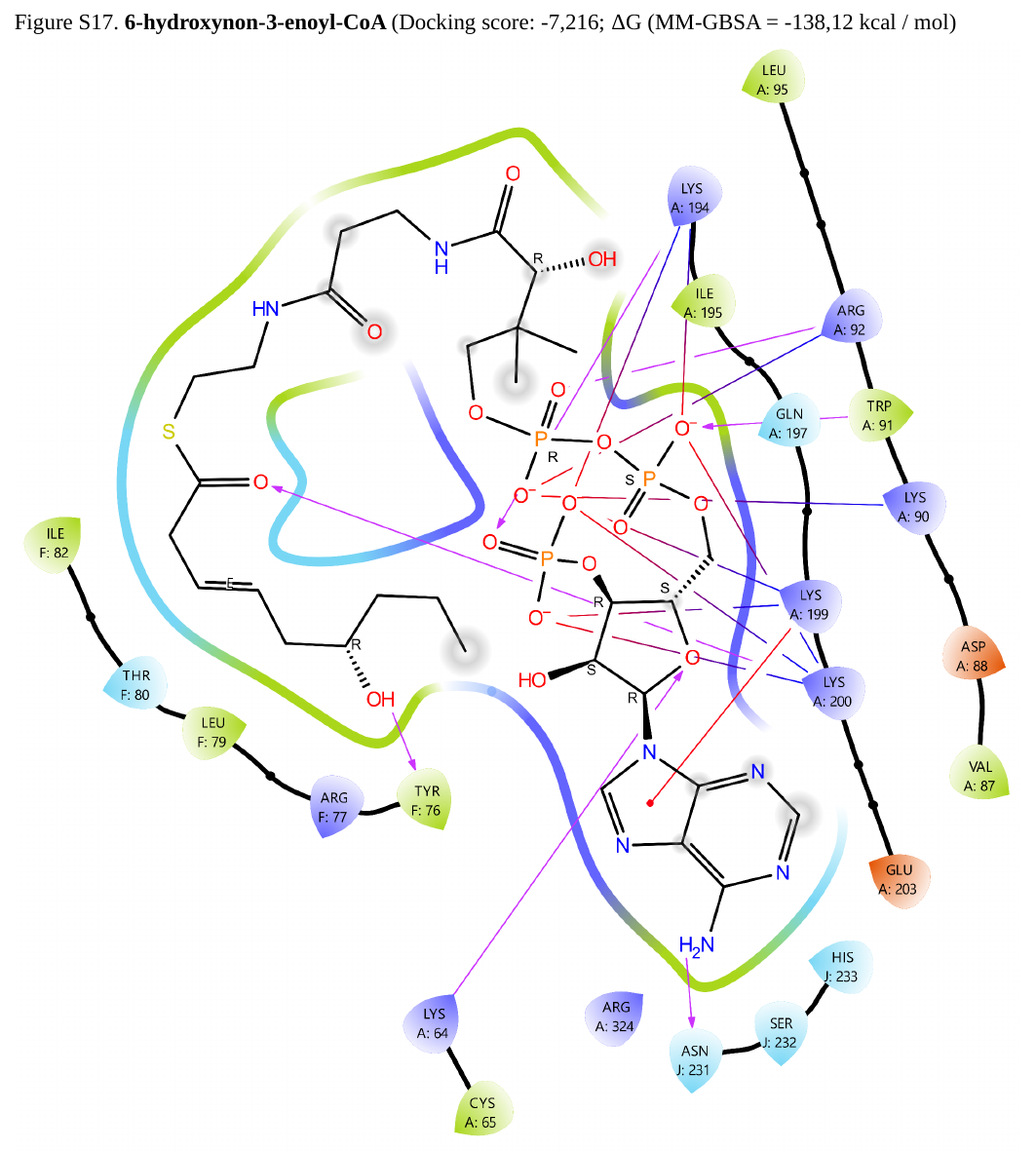

Figure S17. 6-hydroxynon-3-enoyl-CoA (Docking score: -7,216; ΔG (MM-GBSA = -138,12 kcal / mol)

### Slide 19
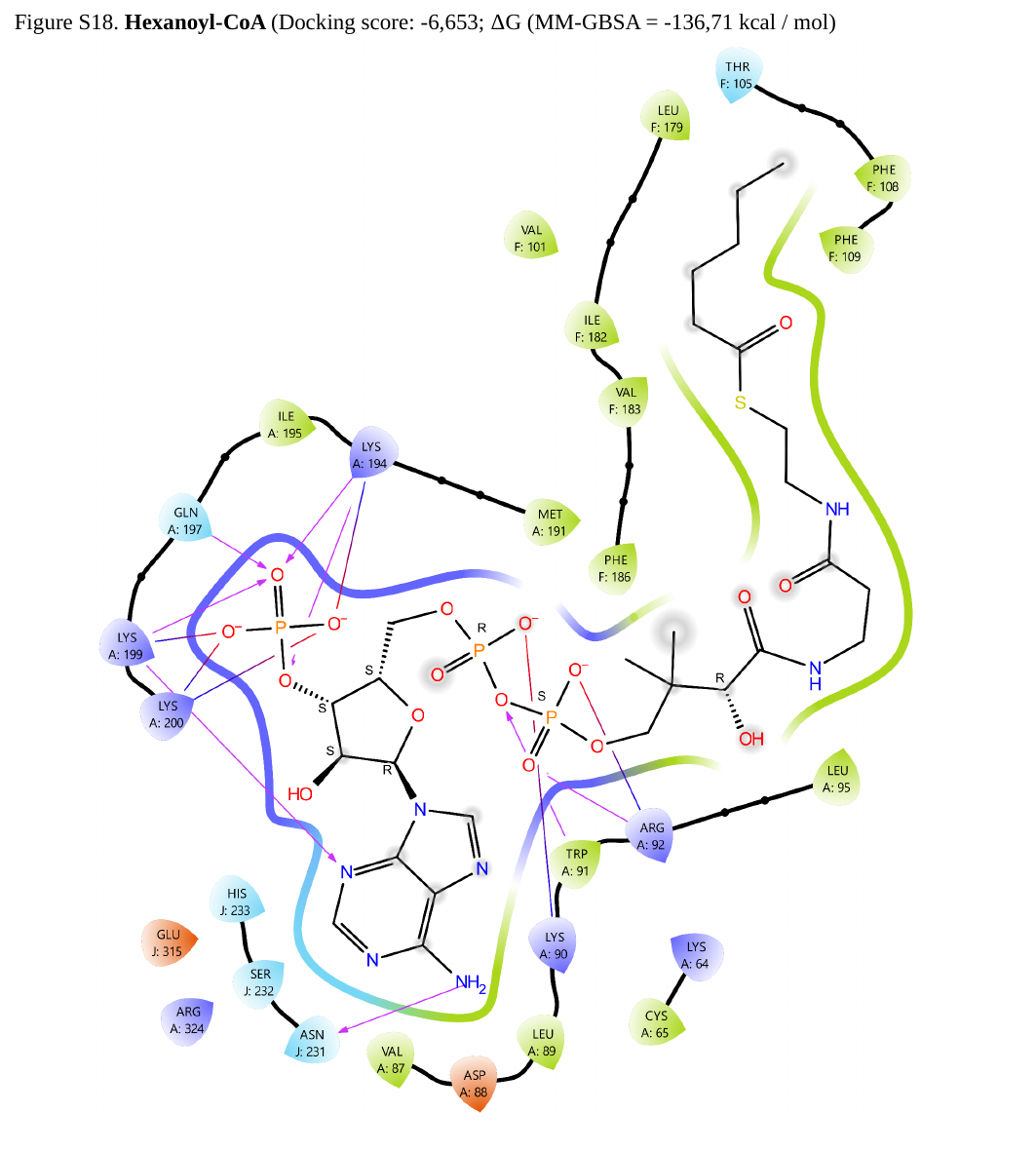

Figure S18. Hexanoyl-CoA (Docking score: -6,653; ΔG (MM-GBSA = -136,71 kcal / mol)

### Slide 20
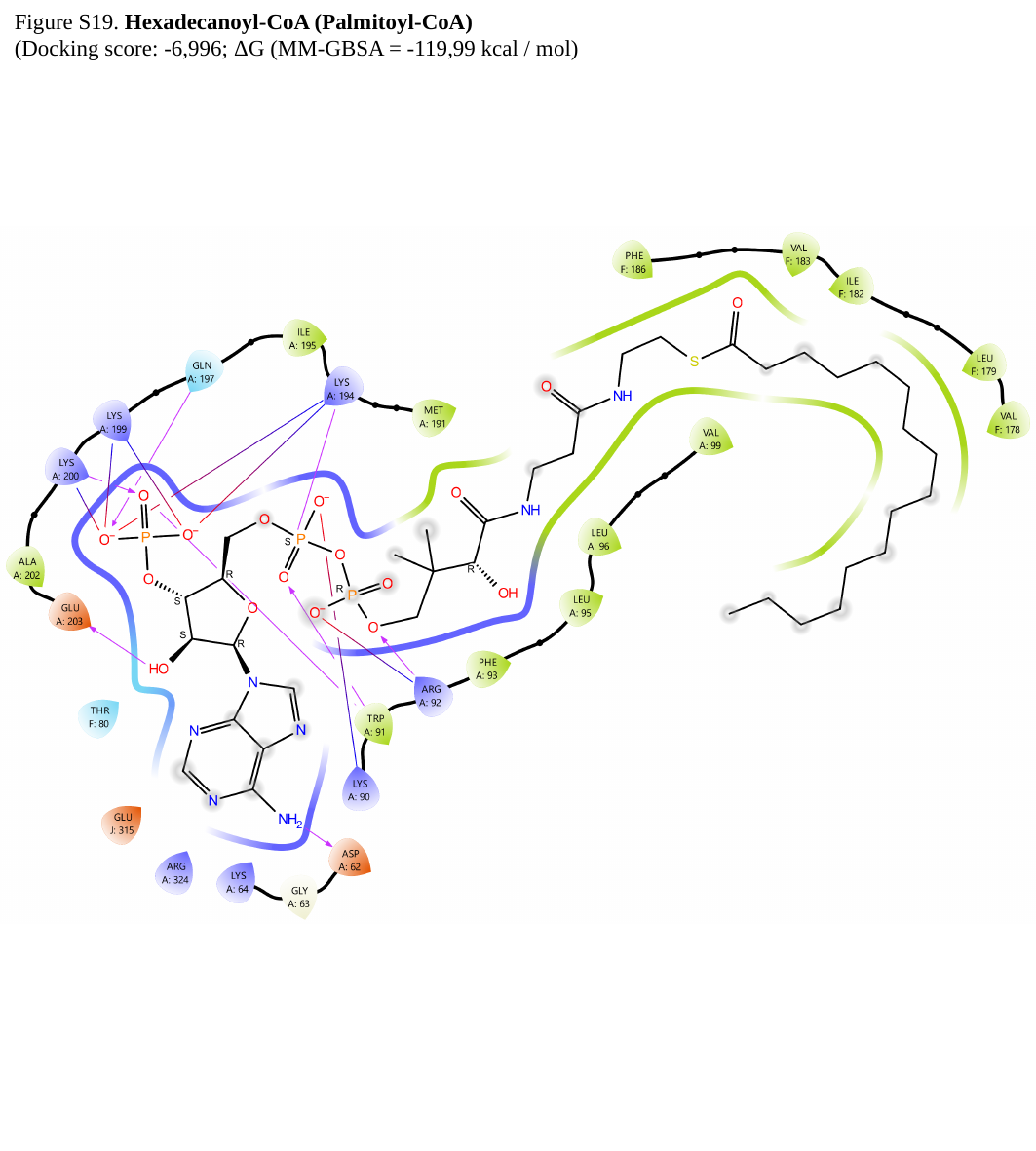

Figure S19. Hexadecanoyl-CoA (Palmitoyl-CoA)
(Docking score: -6,996; ΔG (MM-GBSA = -119,99 kcal / mol)

### Slide 21
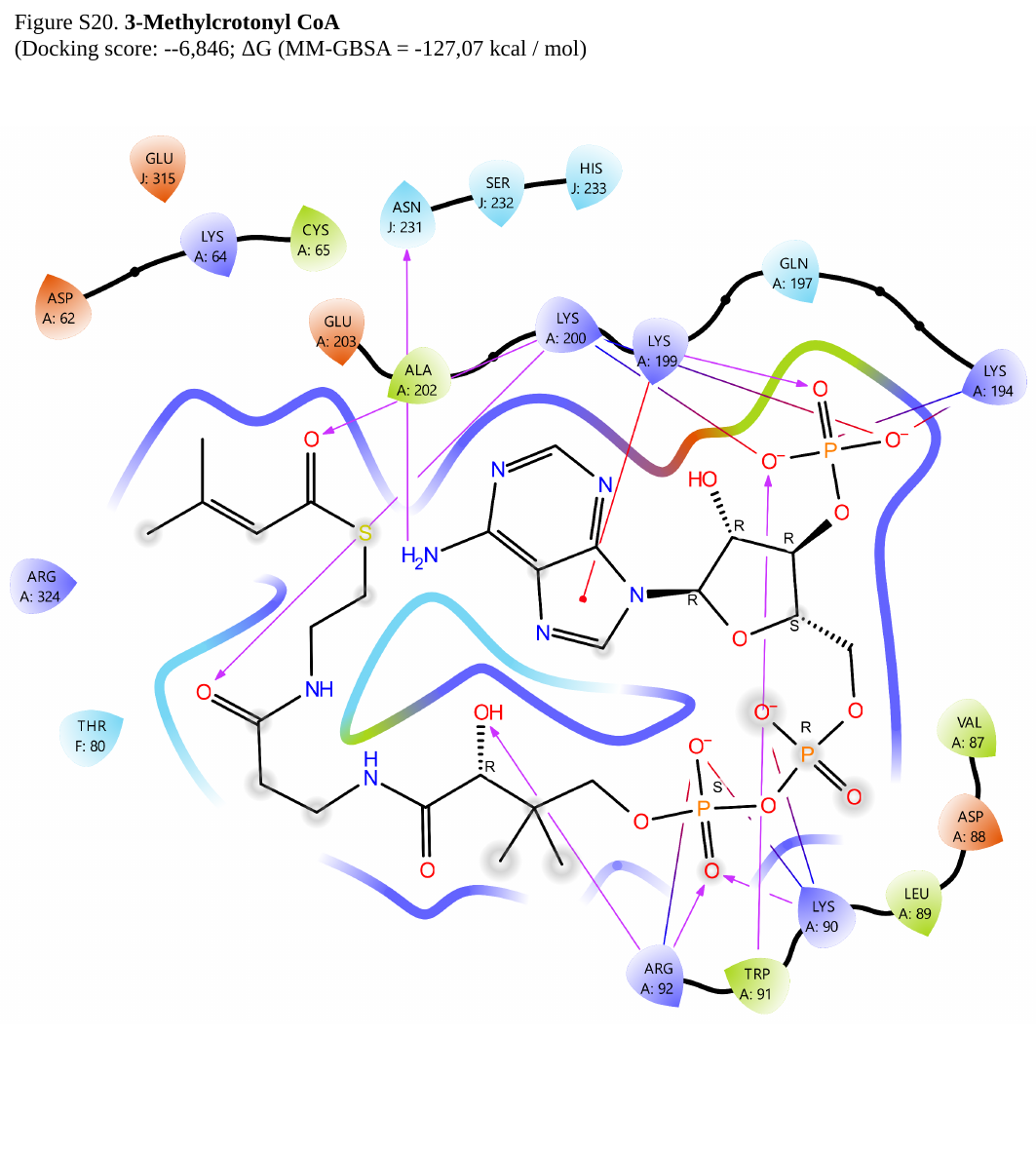

Figure S20. 3-Methylcrotonyl CoA
(Docking score: --6,846; ΔG (MM-GBSA = -127,07 kcal / mol)

### Slide 22
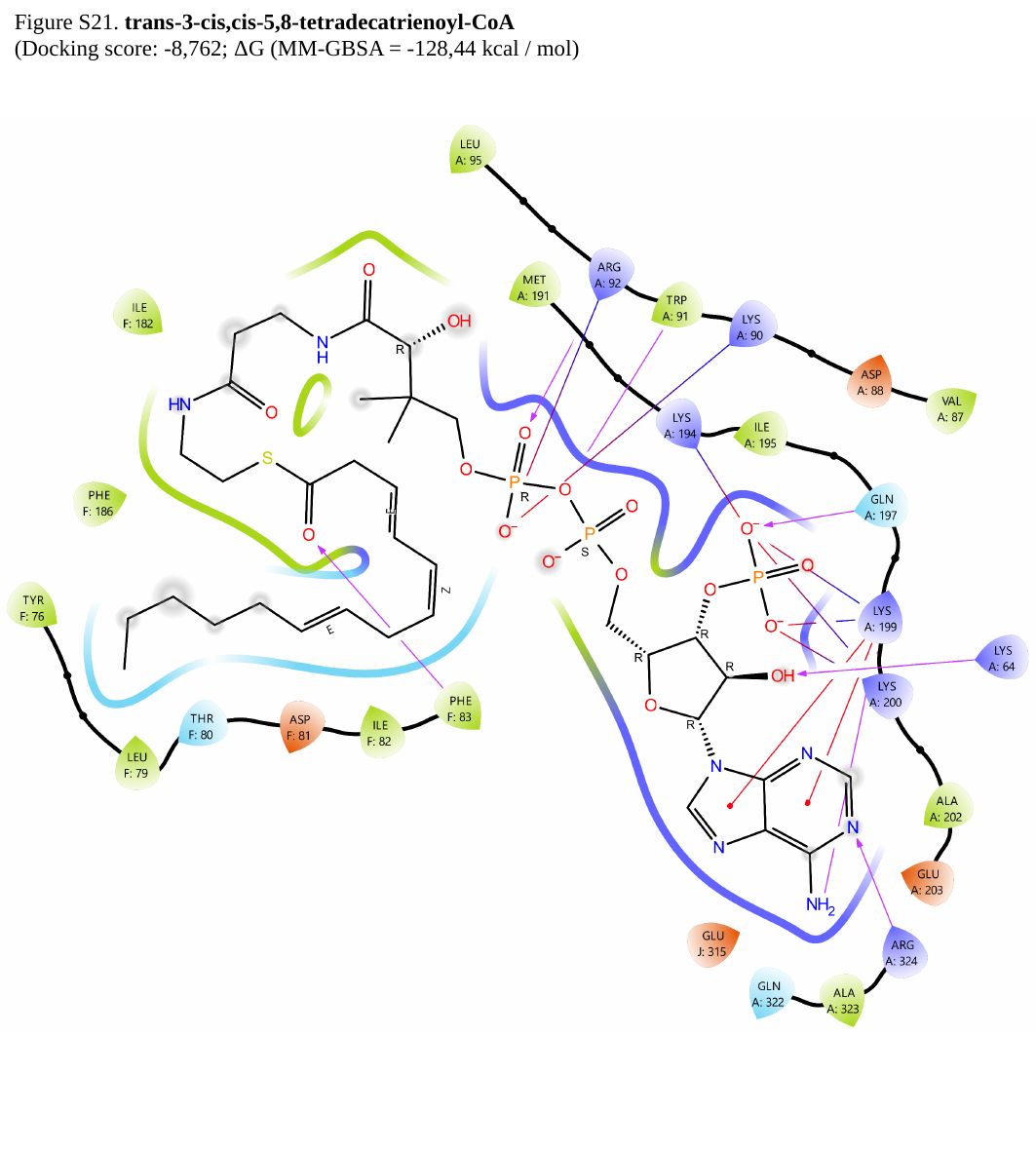

Figure S21. trans-3-cis,cis-5,8-tetradecatrienoyl-CoA
(Docking score: -8,762; ΔG (MM-GBSA = -128,44 kcal / mol)

### Slide 23
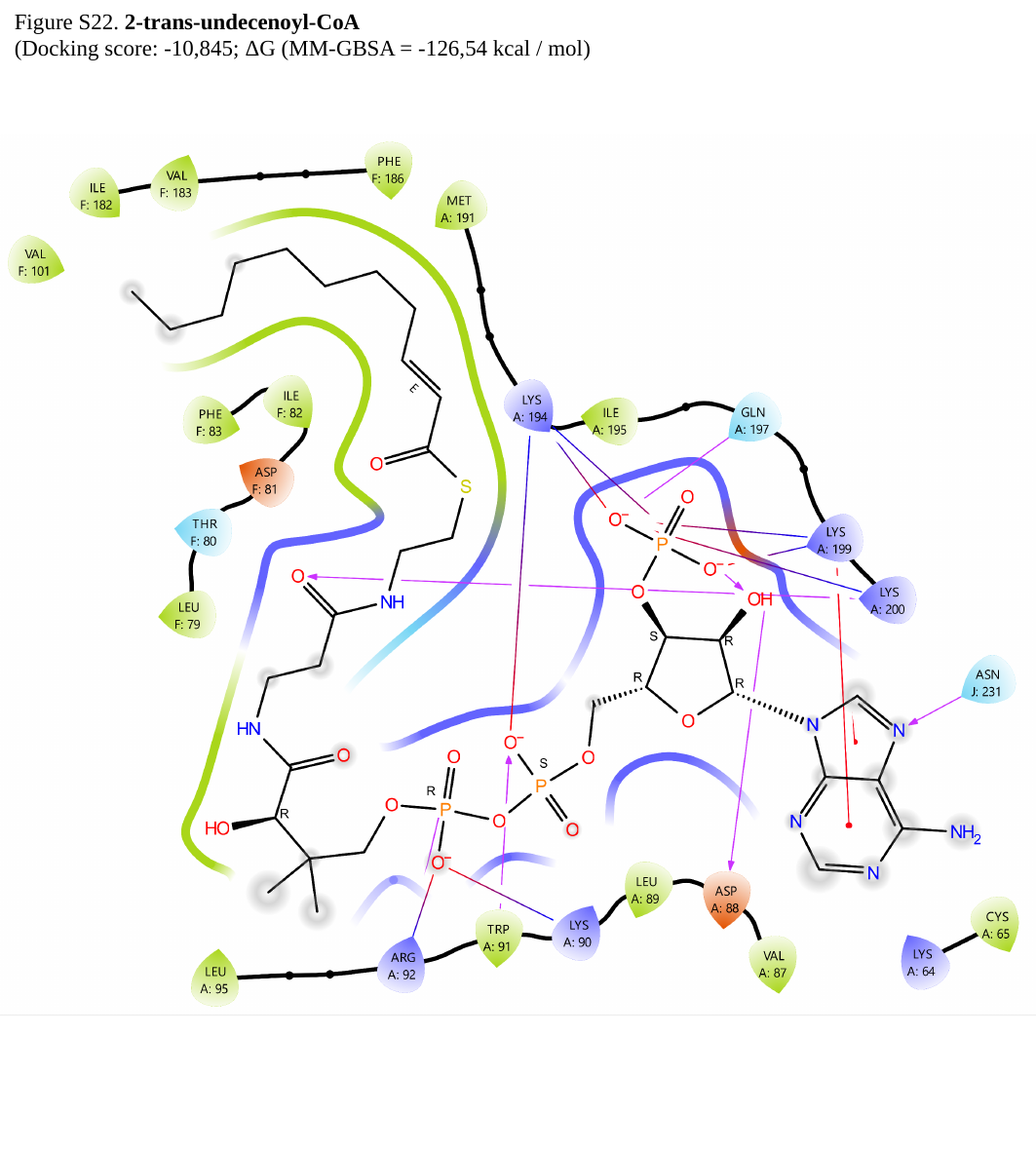

Figure S22. 2-trans-undecenoyl-CoA
(Docking score: -10,845; ΔG (MM-GBSA = -126,54 kcal / mol)

### Slide 24
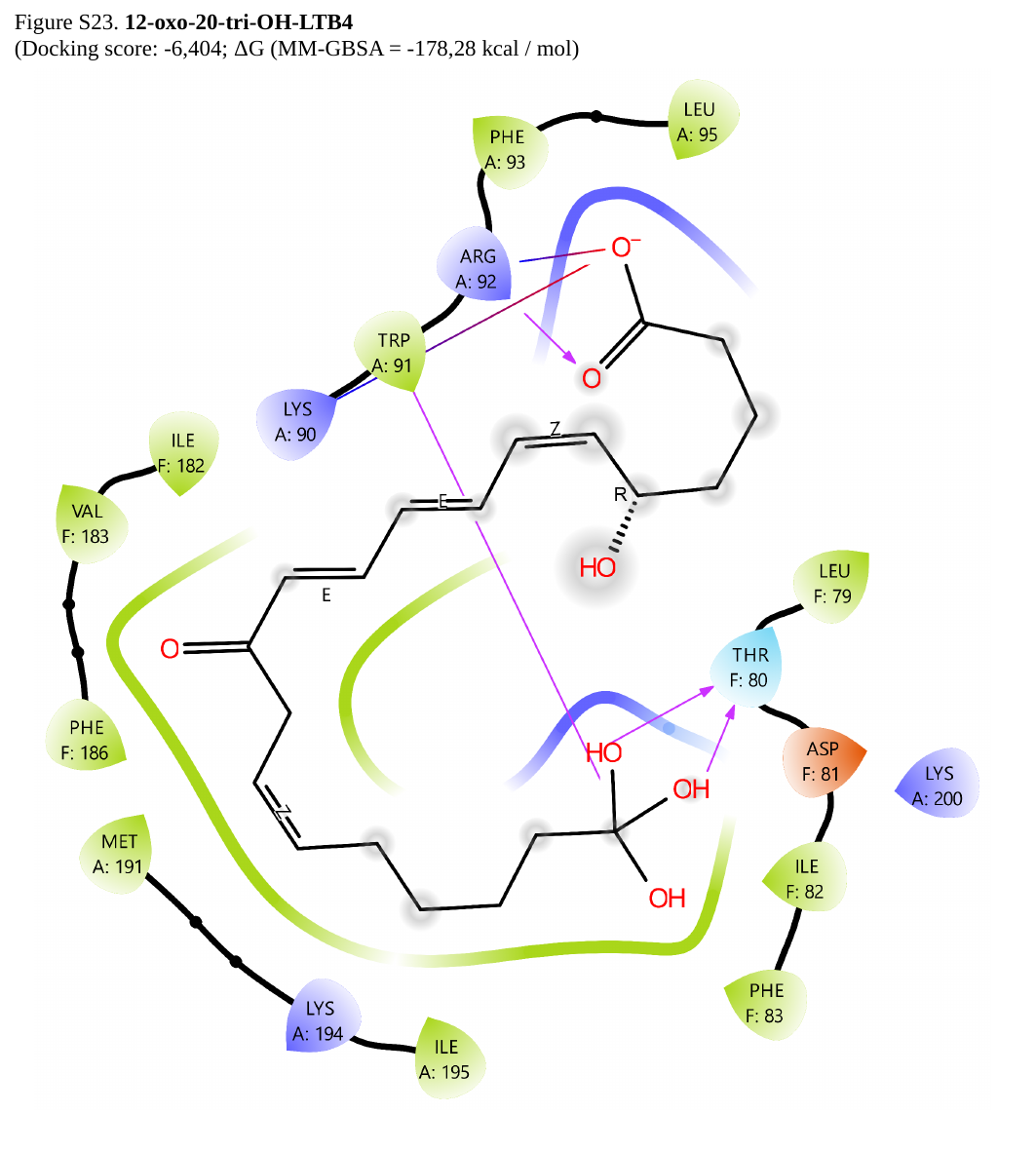

Figure S23. 12-oxo-20-tri-OH-LTB4
(Docking score: -6,404; ΔG (MM-GBSA = -178,28 kcal / mol)

### Slide 25
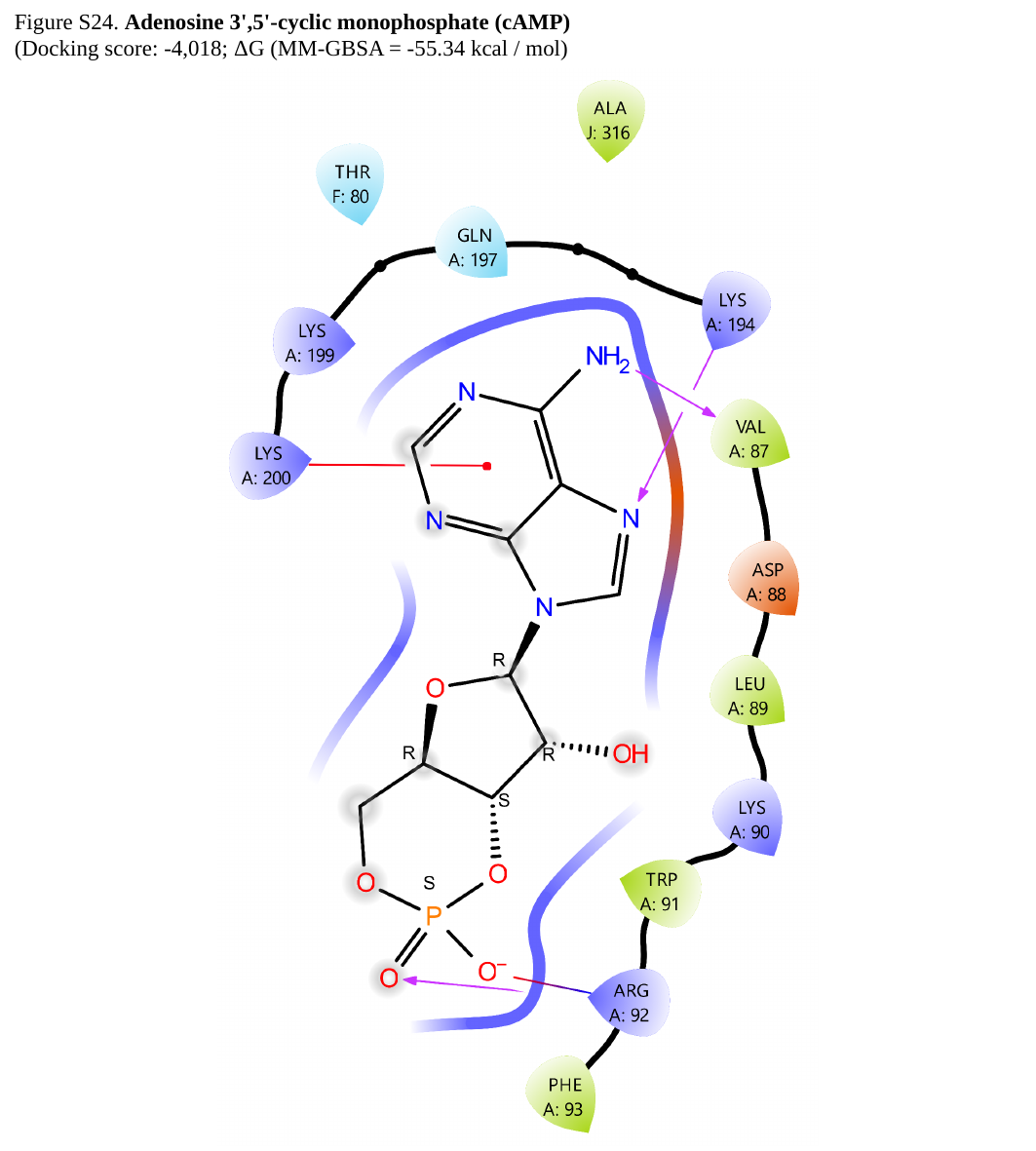

Figure S24. Adenosine 3',5'-cyclic monophosphate (cAMP)
(Docking score: -4,018; ΔG (MM-GBSA = -55.34 kcal / mol)

### Slide 26
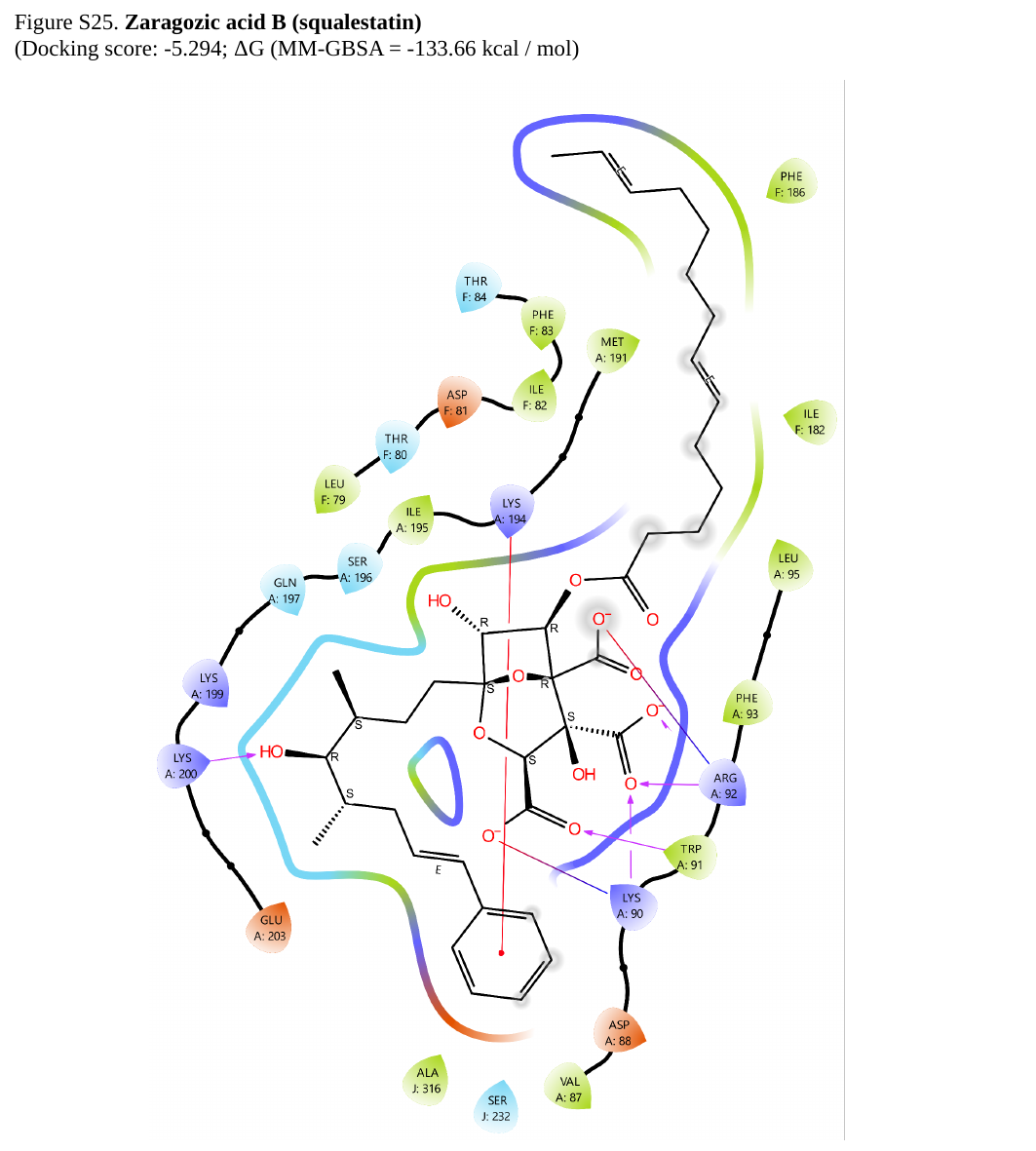

Figure S25. Zaragozic acid B (squalestatin)
(Docking score: -5.294; ΔG (MM-GBSA = -133.66 kcal / mol)

### Slide 27
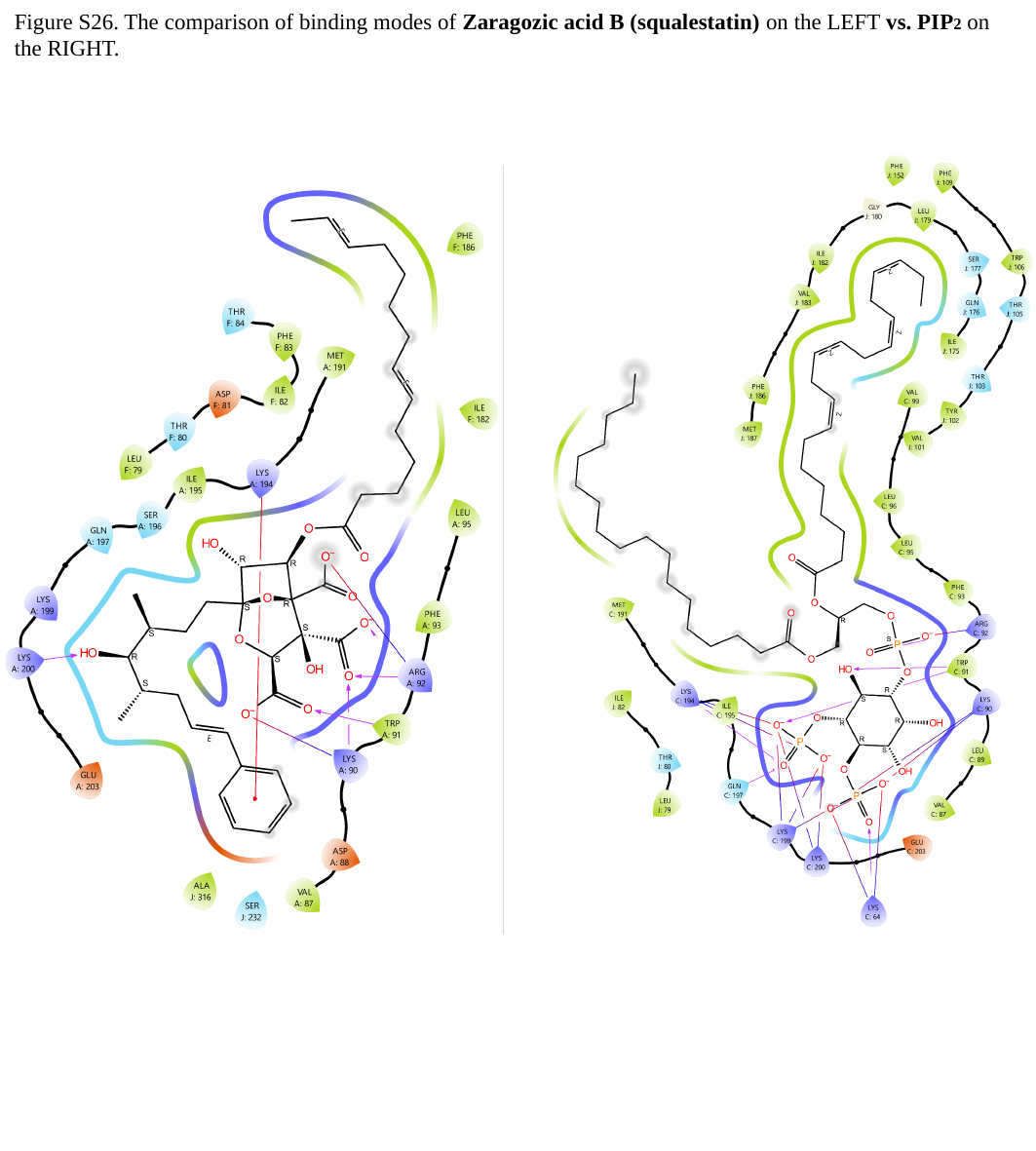

Figure S26. The comparison of binding modes of Zaragozic acid B (squalestatin) on the LEFT vs. PIP2 on the RIGHT.
